# Selection of rAAV Vectors that Cross the Human Blood-Brain Barrier and Target the Central Nervous System Using a Transwell Model

**DOI:** 10.1101/2022.07.06.498990

**Authors:** Ren Song, Katja Pekrun, Themasap A. Khan, Feijie Zhang, Sergiu P. Pasca, Mark A. Kay

## Abstract

A limitation for rAAV-mediated gene transfer into the central nervous system (CNS) is the low penetration of vectors across the human blood-brain barrier (BBB). High doses of intravenously-delivered vector are required to reach the CNS, which has resulted in varying adverse effects. Moreover, selective transduction of various cell types might be important depending on the disorder being treated. To enhance BBB penetration and improve CNS cell selectivity, we screened an AAV capsid-shuffled library using an *in vitro* transwell BBB system with separate layers of human endothelial cells, primary astrocytes and/or hiPSC-derived cortical neurons. After multiple passages through the transwell, we identified chimeric AAV capsids with enhanced penetration and improved transduction of astrocytes and/or neurons compared to wild-type capsids. We identified the amino acid (aa) from region 451-470 of AAV2 associated with the capsids selected for neurons, and a combination of aa from regions 413-496 of AAV-rh10 and 538-598 of AAV3B/LK03 associated with capsids selected for astrocytes. An siRNA screen identified several genes that affect transcytosis of AAV across the BBB. Our work supports the use of a human transwell system for selecting enhanced AAV capsids targeting the CNS and may allow for unraveling the underlying molecular mechanisms of BBB penetration.

## Introduction

Recombinant adeno-associated viral (rAAV) vectors have become one of the most popular gene therapy delivery tools for the clinical treatment of genetic disorders. Zolgensma is an FDA approved gene therapy product which employs an rAAV9 vector to deliver the SMN1 gene into motor neurons for the treatment of patients with lethal spinal muscular atrophy.^1^ Both AAV9 and AAV-rh10 are the current “gold standard” capsids known to cross the blood-brain barrier (BBB) and most often used for delivery into the CNS.^2, 3^ Nonetheless, AAV9 and AAV-rh10 still have limited BBB penetration and CNS transduction. Due to these limitations, high doses of AAV based therapeutics are required to reach the target cells, while a large amount of the vector are taken up by other tissues leading to varying degrees of adverse effects, including death.^4^ Therefore, efficient delivery of cell-type specific rAAV vectors across the BBB represents an important goal for the field.

The BBB is mainly composed of endothelial cells that line the walls of the brain capillaries, which account for 85% of vessel length in the brain.^5^ Tight junctions form between these endothelial cells limit paracellular transport of molecules between the blood and the brain. This ensures that molecular exchange between the circulating blood and the brain is tightly regulated in order to achieve a homeostatic supply of nutrients to the brain, while also fending off harmful molecules to prevent unwanted neuronal excitation. On the neural tissue side of the BBB, pericytes and astrocytes directly associate with the endothelial cells to further strengthen the integrity and functional maintenance of the BBB. Delivery across the BBB remains one of the most efficient way to achieve widespread disseminated gene transfer into the cells in the CNS.^6^

A number of labs have been working towards developing rAAV vectors that can efficiently cross the BBB in rodent or nonhuman primate models (NHP).^7^ However, one of the major limitations is the lack of predictability of AAV transduction efficiency between species. A notable example is AAV-PHP.B, which was selected from a screen in C57BL/6 mice and showed significant improvement over AAV9.^8^ However, this improvement was not observed when AAV-PHP.B was tested in the NHP models.^9, 10^ Further investigations found AAV-PHP.B specifically engages the LY6a receptor that is present only in certain strains of mice^11, 12^. Another challenge is that capsids selected for improved BBB penetration do not necessarily transduce the target cell types with high efficiency. Many rAAV vectors that are able to successfully pass the BBB have a low preference towards neurons, while transduction of supporting cells that directly associate with the BBB, such as astrocytes, is relatively more efficient.^2, 13^ Methods in using transgenic rodent models carrying various cell-type-specific promoters have facilitated the selection of AAVs that specialize in transduction of certain cells.^14^ However, these selection methods remain difficult to adapt to human models. More recently, parallel screens of AAV capsids have been performed in various hiPSC-derived neural cell types, but the BBB penetration abilities of these AAVs remain untested.^15^

To overcome the unpredictable differences in the BBB between species and select for cell type specific AAVs, we have employed an *in vitro* model of the human BBB that consists of a co-culture of human cerebral microvasculature endothelial cells (hCMEC/D3) and primary astrocytes in a transwell system.^16^ We also used hiPSC neurons derived in 3D cortical organoids to select for enhanced transduction efficiency.^17^ We validated this BBB model by demonstrating that AAV9 and AAV.rh10 capsids pass the BBB more efficiently than other wild-type capsid AAVs. Using a barcoded and capsid-shuffled AAV library ^18^ to select for improved capsids in this human BBB transwell system, we were able to isolate a repertoire of capsids with enhanced ability to penetrate the BBB and transduce hiPSC-derived cortical neurons and/or primary astrocytes. These novel AAV capsids not only provide new development opportunities for human gene therapeutics for the CNS, but also serve as tools to identify molecular mechanisms that facilitate BBB penetration and determine CNS cell type specificity.

## Results

### Establish and validate a transwell model of human BBB for selection of AAVs

To overcome the species and cell-type limits of the current AAV vectors, we decided to select AAV vectors in a transwell culture model of the human BBB that is commonly used to study BBB structure and function.^16^ A key component of the BBB is the layer of endothelial cells that form tight junctions to prevent large molecules crossing from the blood into the brain by paracellular transport.^5^ We cultured a confluent layer of human cerebral microvascular endothelial cells (hCMEC/D3) on the top of the layer of the transwell system (Figure S1A). The transwell consists of a polyester membrane with 0.4 μm pores, which keeps the endothelial cells (10 - 20 μm) on the upper side of the transwell, while allowing AAV (∼20 nm) to cross as long as they are able to penetrate the endothelial cell layer. A culture time of 7 days was sufficient to prevent the diffusion of dextran molecules of 70,000 MW (regularly used to test cell layer integrity) and 2, 000, 000 MW (approximately half the size of AAV) from crossing the transwell (Figure 1A and B). Furthermore, a higher number of AAV9 penetrated the layer of hCMEC/D3 cells into the media below (flowthrough) as compared to AAV2 (Figure S1B). This observation aligns well with previously published data comparing AAV9 and AAV2 in the same transwell model using primary human brain microvascular endothelial cell cultures.^19^ As astrocytes are directly associated with endothelial cells in the BBB, we added a layer of primary human astrocytes to the bottom side of the porous membrane (Figure S1A). Addition of the astrocytes did not interfere with the dextran permeability of the hCEMC/D3 cells (Figure 1A) and allowed us to perform selection of AAVs that would enter the astrocytes after crossing the hCMEC/D3 cells.

**Figure 1.**
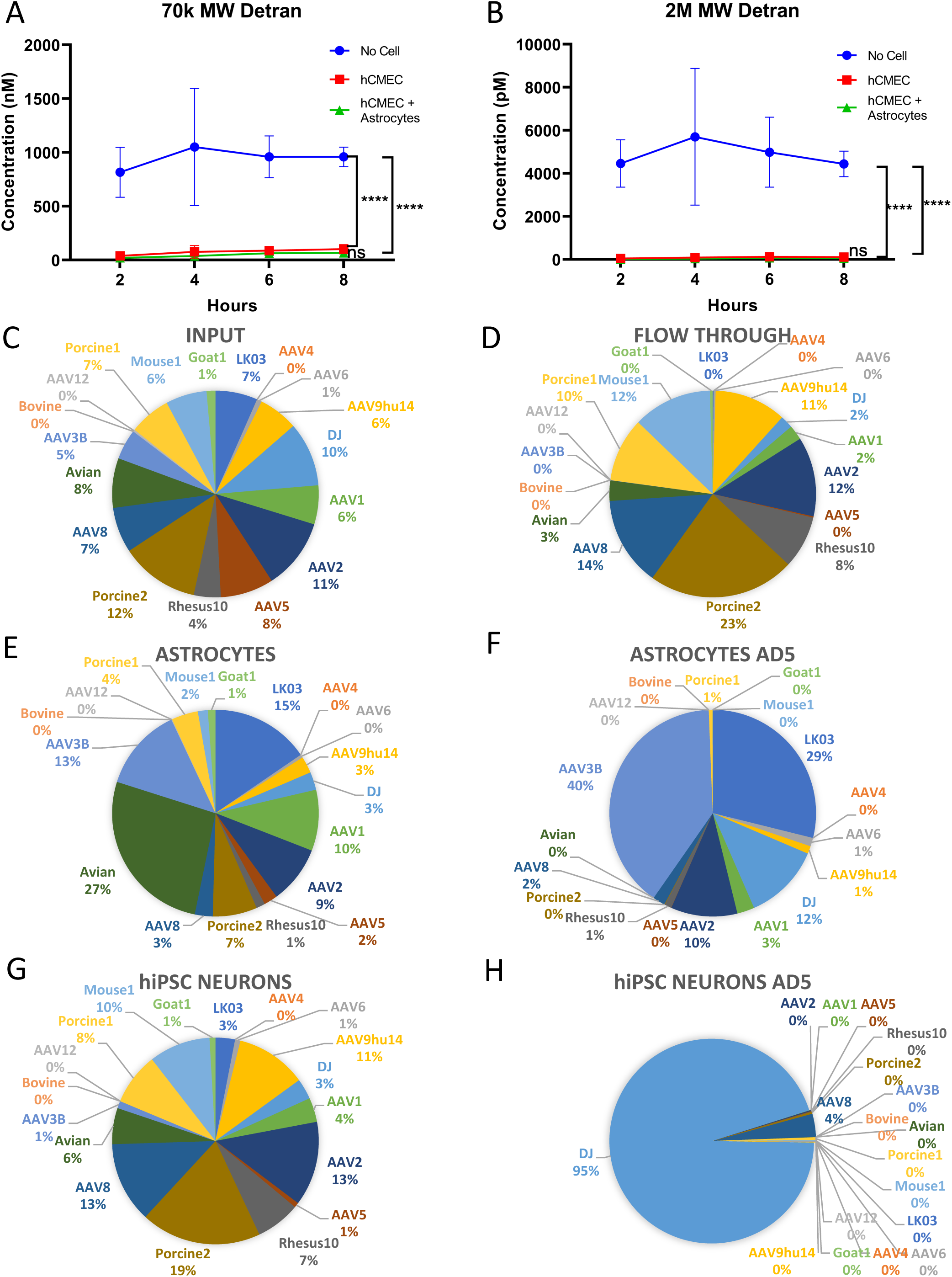
Establish and validate a transwell model of human BBB for selection of AAVs. A. and B. Permeability of transwells with no cells, hCMEC/D3 or hCMEC/D3 and human primary astrocytes were tested using 70k MW (A) and 2M MW (B) dextran. Significance determined by 2-way Anova test, **** p<0.0001. C-H, 18 uniquely barcoded AAVs were pooled and passaged through the transwell BBB model with hCMEC/D3, primary astrocytes and hiPSC-derived cortical neurons with MOI of 1e4. The percentage of each of the 18 AAV is graphed in the input (C), flowthrough media (D), primary astrocytes with (F) or without (E) Ad5, and in hiPSC-derived cortical neurons with (H) or without (G) Ad5.

To further demonstrate that the trans-well system indeed recapitulates the human BBB in allowing only certain known AAV vectors to cross efficiently, we tested a pool of 18 uniquely barcoded AAVs by passaging them through our model system. This pool of barcoded AAVs had been previously developed by our group to test other model systems and to optimize the AAV selection strategy.^18, 20^ The pool included AAVs known to cross the BBB more efficiently, such as AAV9 and AAV-rh10, and those previously described not to cross the BBB (e.g AAV-DJ).^2, 3^<u>_ENREF_3</u> When tested in the transwell BBB model, we first compared the virus composition in the input and flowthrough of the transwell and found that AAV-rh10 and AAV9 are among the viruses whose proportions increased in the flowthrough compared to input (Figure 1C, 1D, S2A and S2B). Depending on the MOI used, the increase was 2.0- to 2.8- fold for AAV-rh10 and 1.8- to 1.9- fold for AAV9 (Supplemental Table1). AAV8, AAV-Mouse1, AAV-Porcine1 and AAV-Porcine2 also showed increased proportions of up to 2.0- fold in the flowthrough (Supplemental Table1). Comparatively, flowthrough proportions of AAV-DJ decreased more than 4-fold in comparison to the input. The flowthrough from each transwell was also added to hiPSC-derived neurons, isolated by immunopanning from human cortical organoids, to evaluate the transduction efficiency of the respective AAVs (Figure S1A). We found that the proportion of viruses that transduced the hiPSC-derived cortical neurons were not significantly different from that in the flowthrough (Figure 1D, 1G, S2A, S2B and Supplemental Table1). These results suggest that the pool of AAVs found in the flowthrough was a good indicator for neuron transduction efficiency.

However, the composition of the pool of viruses that entered the layer of primary human astrocytes was drastically different from that of the flowthrough and in hiPSC-derived neurons (Figure 1E, S2A and S2B). The AAVs that showed a consistent increase in astrocytes across of all MOIs tested include AAV1 (1.2- to 1.9- fold), AAV3B (1.5- to 2.8- fold), AAV-LK03 (1.8- to 2.5- fold) and AAV.Avian (2.2- to 3.4- fold) (Supplemental Table1). These data suggest that the transwell BBB model will allow us to select for BBB-penetrating capsids with different preferences in transducing different types of cells in the CNS.

We then proceeded to evaluate how the addition of Adenovirus 5 (Ad5) to the hiPSC-derived cortical neurons or primary astrocytes would affect the proportion of AAVs that transduced these cells. We found that Ad5 resulted in the dominance of variants that replicate well in the respective cell type, a phenomenon that had been previously described.^20^ We found that AAV-DJ or AAV2 had become the dominating species in hiPSC-derived cortical neurons (Figure 1H, S2A and S2B), while AAV3B and AAV-LK03 had taken over more than 68% of the population in astrocytes (Figure 1F, S2A and S2B). Although removing Ad5 from the selection process may alleviate this bias, the capsids selected in the absence of Ad5 may not necessarily exhibit improved transduction efficiency since multiple steps after cell entry are important for successful transgene expression. In addition, we observed that viral copy numbers were very low when Ad5 was omitted from the screen, making consecutive rounds of selection difficult. Thus, we decided to conduct hybrid schemes of selection of our AAV library with or without addition of Ad5.

### Selection of AAV vectors that efficiently cross the BBB and transduce hiPSC-derived cortical neurons and primary astrocytes

Over the past 20 years, our lab has established methods to make complex multispecies interbred AAV capsid libraries.^18, 21–24^ More recently, we have generated capsid-shuffled libraries where each AAV harbors a short and unique barcode sequence downstream of the *cap* polyA signal.^18, 25^<u>_ENREF_17</u> This method allowed us to track AAV variant enrichment by high-throughput sequencing of the barcodes. We added an AAV library that contained chimeric capsids derived from 10 parents to the top of the transwell human BBB model and performed three separate selection experiments (Figure 2). The selections were carried out in two consecutive rounds in the flowthrough, hiPSC-derived cortical neurons and primary astrocytes. For the selection in flowthrough that enriched for AAV variants capable of BBB penetration, we did not use Ad5 to replicate viral variants throughout the entire process. We also did not include Ad5 during the selection process of capsids that efficiently cross the endothelial cell layer and transduce neurons or astrocytes. However, to generate sufficient input for the second round of selection, Ad5 was added to the neurons and astrocytes after the first round of selection. The barcode sequences of AAV from the input and output of each round of the three selections were identified by Illumina sequencing. Each selection was done in biological replicates and we identified barcodes that were present in both replicates and were continuously enriched in two rounds of selection. We found a total of 25, 90 and 250 such barcodes from the flowthrough, neuron and astrocyte screens, respectively.

**Figure 2.**
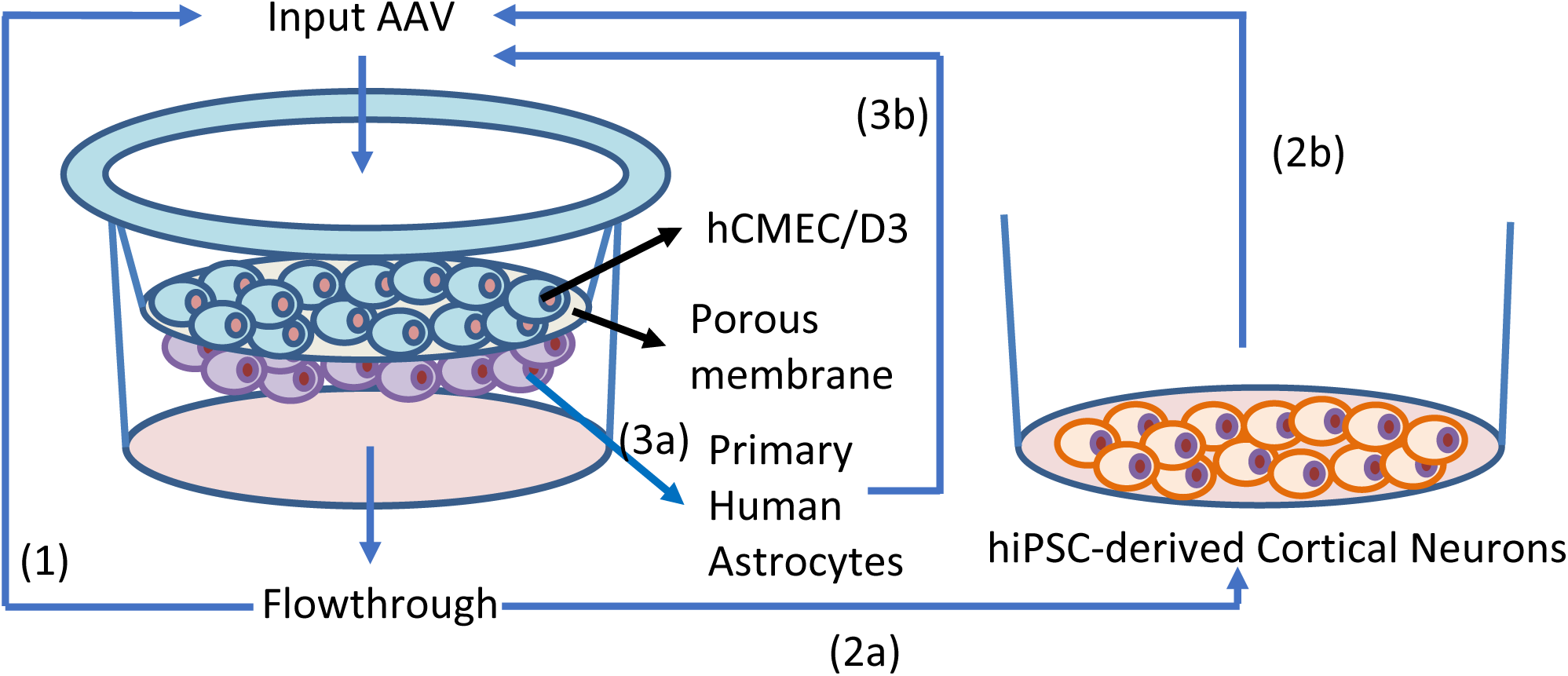
Selection of AAV vectors in the human BBB transwell model. Three separate selection schemes were conducted using an barcoded and capsid-shuffled AAV library (MOI 1e6) as input added to the top of the transwell containing hCMEC/D3 and primary astrocytes. (1) AAVs collected from media in the Flowthough at 24h were determined by sequencing and were added to a new transwell for a second round of enrichment. (2a) Media from Flowthrough at 24h was added to hiPSC-derived cortical neurons to select for AAVs that transduce this cell type. (2b) In a separate culture of hiPSC-derived neurons, media from Flowthrough at 24h was added along with Ad5 to generate input for the second round of selection in transwell and in hiPSC-derived neurons as in (2a). (3a) Astrocytes from the transwell were collected at 24h to determine AAVs that transduce this cell type. (3b) In a separate transwell, Ad5 was added to the astrocytes at 24h to generate input for the second round of selection in astrocytes as in (3a).

### Validation of selected rAAV vectors in crossing the transwell BBB model

We vectorized a total of 14 AAV capsid sequences, 2 from the flowthrough selection, 6 from the neuron selection, and 6 from the astrocyte selection. Each rAAV capsid was used to package a firefly luciferase expression construct under transcriptional control of a CAG promoter (CAG-FLuc). We first tested each capsid variant separately for their ability to cross both the hCMEC/D3 and primary astrocyte cell layers in the transwell BBB model by measuring the abundance of vector genomes in the flowthrough (Figure 3A). The AAVs from the flowthrough and neuron selections crossed the transwell with high efficiency as early as 3 hours post transduction (hpt) and continued to transcytose during the 24-hour testing period. When compared to the control AAV9 at 24hpt, we found 5 of the rAAV vectors were 2.8- (RS.N8d), 8.4- (RS.R3), 5.7- (RS.R6), 14.8- (RS.R11) and 8.9- fold (RS.R18) more efficient. Two of the rAAV vectors, RS.R4 and RS.R5, showed similar efficiency to the controls. RS.N2 was the only variant that was less efficient at crossing than AAV9 and AAV-rh10. The AAV capsids selected from the astrocyte screen, on the other hand, were all less efficient at crossing the hCMEC/D3 and primary astrocyte cell layers especially at 3 and 6 hours post transduction. Although RS.A5a1 and RS.A5a2 reached similar levels to the two control AAVs at 24hpt, the four other AAVs remained less efficient when compared to AAV9 and AAV.rh10.

**Figure 3.**
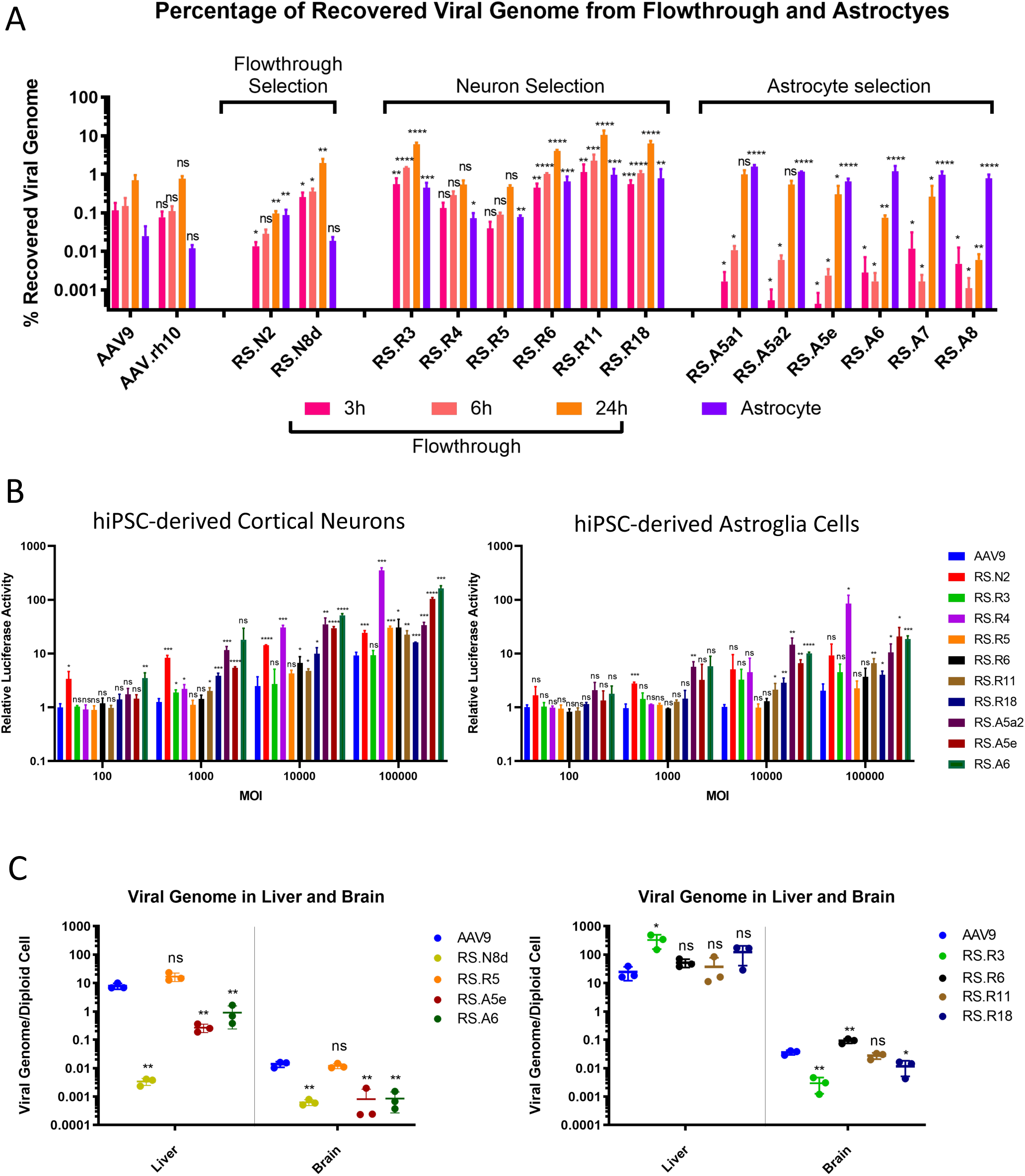
Validation of Selected rAAV vectors in transwell, hiPSC-derived cortical neurons and astroglia cells, and mice. A. 14 AAV capsids from the three selections were vectorized and tested in the transwell BBB system with hCMEC/D3 and primary astrocytes, each rAAV was tested in separate transwells and viral genome copy numbers as a percentage of input virus were determined in flowthrough media at 3, 6 and 24h and in astrocytes at 24h. B. Transduction efficiency 12 AAV capsids from the three selections packaged with firefly luciferase expression cassette were tested in hiPSC-derived cortical neurons and astroglia cells. Relative luciferase activity as compared to AAV9 in each cell type were determined after 72h. C. Delivery of viral genomes to mouse liver and brain after retro-orbital injection was determined for RS.N8d, RS.R5, RS.A5e and RS.A6 at 21 days and for RS.R3, RS.R6, RS.R11 and RS.R18 at 28 days. N >= 3 for all conditions, significance determined by Student’s t test; ns, not significant; * p<0.05; ** p <0.01; *** p<0.001; **** p<0.0001.

We selected the representative rAAV vectors from each selection (RS.R5, RS.R6, RS.R11, RS.R18, RS.A5e and RS.A6), and package each capsid with a CAG human frataxin (CAG-hFXN) expression construct with a unique barcode sequence between the stop codon and polyA signal. This allowed us to pool the AAV capsids together with AAV9 and AAV-DJ at equimolar ratios, and tested their ability to cross the same transwell BBB when subjected to competition (Figure S3B). We found that the data from this pooled approach correlated well with the data obtained when each virus was tested separately. RS.R6, RS.R11 and RS.R18 remained the most efficient variants at crossing into flowthrough, RS.R5 performed similar to AAV9, while RS.A5e, RS.A6 and DJ were the least efficient capsid variants. Taken together, these data suggest that our flowthrough and neuron selection schemes in the transwell BBB model were highly effective at enriching for AAVs that cross the endothelial and astrocyte layers efficiently.

### Validation of selected rAAV vectors in transducing primary astrocytes in the transwell BBB model

Using the same transwell experiments as described above, we examined the efficiency of each rAAV capsid variant in transducing the primary astrocytes after crossing the endothelial cells. We found that all rAAV variants from the astrocyte selection delivered higher amounts of vector genome DNA to astrocytes than the two control serotypes AAV9 and AAV-rh10 (Figure 3A). These rAAV vectors also delivered higher proportions of vector DNA to astrocytes compared to the flowthrough. Apart from RS.N8d, the rAAV capsids derived from the flowthrough and neuron screens were also capable of delivering higher amounts of vector DNA to astrocytes when compared to the control AAVs. However, in contrast to the variants derived from the astrocyte screen, delivery of vector genomes into astrocytes was less efficient than the flowthrough media. When we tested the rAAV vectors in the pooled approach, we found that RS.A5e and RS.A6, two capsid variants enriched in the astrocyte selection, were the most effective at delivering vector genome DNA to astrocytes (Figure S3C). When expression of vector RNA was tested, RS.A6 remained the most dominant in astrocytes, while the performance of RS.A5e dropped behind RS.R11 and RS.R18, but still remained more efficient than the other vectors in the pool (Figure S3C). These data suggest that our astrocyte selection scheme in the transwell BBB model can select for AAV capsid variants that cross the endothelial layer and transduce astrocytes.

### Testing transduction efficiency of the selected rAAV vectors in hiPSC-derived cortical neurons and astroglia cells as well as hCMEC/D3

The transduction efficiency of the selected vectors was examined in hiPSC-derived neurons and astroglia that were isolated from cortical organoids using immunopanning for Thy1 and HepaCAM as previously shown. ^26^ We transduced the two cell types with rAAV vectors carrying the CAG-FLuc construct at increasing multiplicity of infection (MOI) from 100 to 100,000 (Figure 3B). Apart from RS.R3, all rAAV vectors from the flowthrough and neuron selection were more efficient at transducing hiPSC-derived neurons than AAV9, especially at a higher MOI. Notably, while RS.N2 and RS.R4 were the most efficient at direct transduction of hiPSC-derived neurons, these two rAAV vectors were the least efficient at penetrating the endothelial and primary astrocyte cell layers (Figure 3A), suggesting a negative correlation between the ability to transduce and the efficiency to cross the BBB in this model system. RS.N2, RS.R4, RS.11 and RS.18 were also more efficient at transducing hiPSC-derived astroglia at a higher MOI. All three capsid derived from the astrocyte screen were more efficient at transducing both human cortical organoid neurons and astroglia as compared to AAV9. We also examined the transduction of hCMEC/D3 cells with the pooled rAAV vectors RS.R5, RS.R6, RS.R11, RS.R18, RS.A5e, RS.A6, AAV9 and AAV-DJ carrying uniquely barcoded CAG-hFXN construct (Figure S3E and S3F). While AAV-DJ was the dominating species both in terms of vector DNA delivery and RNA expression, all representative rAAV vectors derived from our various screens also showed higher transduction of endothelial cells when compared to AAV9.

### Evaluation of the selected rAAV vectors in a mouse model

Although mouse models may not be the most relevant for the prediction of AAV capsids that can cross the human BBB, we examined representative rAAVs selected in our screens for their ability to transduce the liver and brain in comparison to AAV9 (Figure 3C). We found that RS.R6 was 2.6- fold better than AAV9 at transducing the mouse brain, while transduction efficiency for liver was similar between the two capsids. RS.R3 was more efficient at transducing liver but less efficient at targeting the brain. For all other rAAV variants, although in our *in vitro* human BBB transwell model they were predicted to be superior to AAV9, in the mouse model they were either similar or less efficient at transducing both the brain and liver. These findings further suggest that mouse models may not be the most relevant for evaluating AAVs in crossing the human BBB.

### Xover analysis of AAV capsid sequences reveal specific amino acid patterns that may contribute to BBB crossing phenotype

To understand if the distribution of certain amino acids (aa) contributed to the observed phenotypes of the selected capsid variants, we performed Xover analysis using the protein sequences of each capsid (Figure 4).^27^ This analysis reveals the crossover patterns of each chimeric vector, where the parental sequences aligning to the longest uninterrupted stretch in the shuffled capsid sequence are chosen as the most probable contributing source. By comparing the graphs generated from Xover analysis, we noticed two distinct patterns in the ∼450 to 650 aa region (Figure 4).

**Figure 4.**
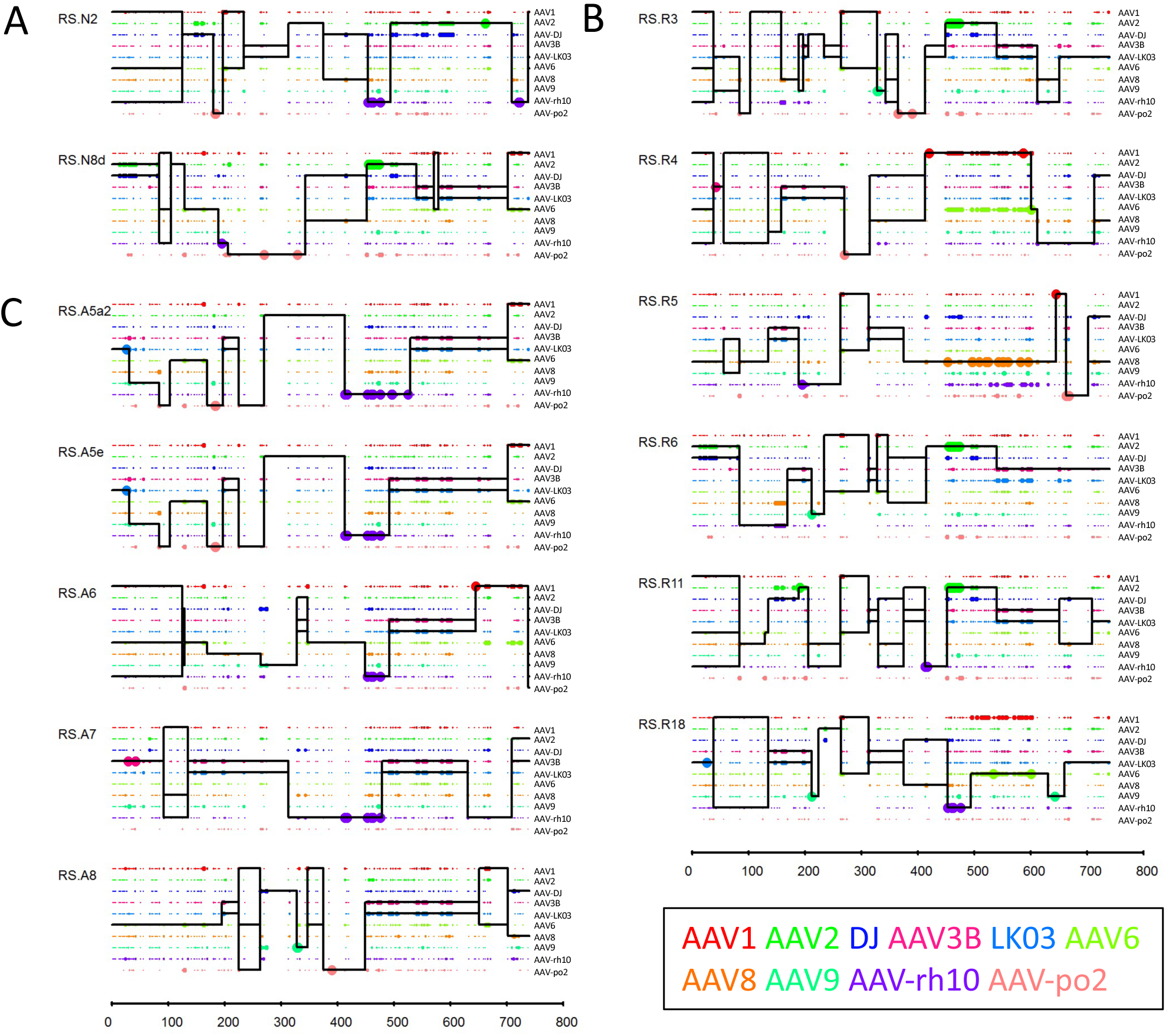
Xover analysis of vectorized AAV capsid amino acid sequences. Xover analysis were performed for each of the vectorized AAV capsids from flowthrough (A), neuron (B) and astrocyte (C) selection. RS.A5a1 was excluded as it only has one amino acid (L464Q) difference from RS.A5a2. Parental AAVs are depicted in different colors. Large dots represent 100% match to the parental AAV amino acid and small dots represent more than one parental match at the position. The horizontal black lines represent the AAV parents contributing to each rAAV capsid chimera. The vertical black lines are predicited crossover positions between parents.

Specifically, all AAV capsids from the astrocyte selection with the exception of RS.A8 harbor the stretch between aa 453 to 477 from AAV-rh10 and a stretch between aa 526 to 627 from AAV3B/LK03 (Figure 4C). The 453 to 477 aa region of AAV-rh10 includes three unique amino acids (453S, 459G and 462Q) in the variable region (VR) IV ^28^, and three additional aa (468A, 472N and 475A) in close proximity of VR IV that are suggested to confer the ability to cross the BBB ^29^. All AAV capsids from the astrocyte screen contains the aa 526 to 627 from AAV3B/LK03. This aa region covers many unique amino acids within or in proximity to VR VI (538H and 540N), VII (549T and 554E) and VIII (582N, 592T, 594R and 598D). Interestingly, RS.N2 and RS.R18 which were derived from the flowthrough and neuron selection screens, respectively, also contain AAV-rh10 aa sequence from 453 to 477. However, neither of these capsids share the aa 526 to 627 region of AAV3B/LK03. Thus, the unique combination of AAV-rh10 aa 453 to 477 and AAV3B/LK03 aa 526 to 627 may contribute to the phenotype of efficiently crossing the BBB endothelial cells and transducing astrocytes.

Another pattern that is less prominent is the combination of AAV2 aa 449 to 535 and AAV3B/LK03 aa 537 to 608. Three of the AAV capsids enriched in the neuron screen (RS.R3, RS.R6 and RS.R11) as well as RS.N8d from the flowthrough screen harbor this pattern (Figure 4A and 4B). The aa 449 to 535 region derived from AAV2 encompasses a total of six unique amino acids, four (451P, 456T, 461Q and 467A) within and two (469D and 470I) in close proximity to VR IV. There are also three additional amino acids (492S, 493A and 500Y) that are part of VR V and are shared only between AAV2 and AAV-DJ. Although the AAV3B/LK03 derived aa stretch between 537 and 608 is a shorter region compared to that observed in the AAV capsids selected in the astrocyte screen, both segments encompassed the same unique amino acids. The unique combination of AAV2 aa 449 to 535 and AAV3B/LK03 aa 537 to 608 may contribute to the improved endothelial cell and astrocyte layer penetration and the enhanced transduction of hiPSC-derived cortical neurons in our BBB model system.

### Long read sequencing analysis confirms specific amino acid patterns in AAV capsid sequences from different selections

The aa patterns identified in the of vectorized AAV capsid sequences were intriguing, but the number of sequences analyzed was very limited. This prompted us to examine a broader number of capsid sequences by submitting our round one and round two selections to Pacific Biosciences SMRT single DNA molecule long read sequencing to obtain full length capsid sequences along with its barcode.^20^ From the 25, 90 and 250 barcodes that were continuously enriched in two rounds of duplicated selection in the flowthrough, neuron and astrocyte, we were able to pull out 6, 28 and 43 associated full capsid sequences, respectively. For every identified AAV capsid sequence, we calculated the probability of the contribution of each parental AAV at every amino acid position. We then averaged the parent contribution at each amino acid position for all 6, 28 and 43 capsid sequences from the flowthrough, neuron and astrocyte screens, respectively, and calculated fold change and statistical significance comparing to the average of 11613 sequences from the unselected input AAV library.

For the capsids from the flowthrough selection screen, four aa from AAV2 (451P, 456T, 461Q and 469D) were identified as significantly enriched (Figure 5A, S4A, Supplemental table 2). The enriched aa in capsids from the neuron selection screen share the same four aa, with two additional (467A and 470D) also from AAV2 (Figure 5B, S4B, Supplemental table 2). These positions were also identified in the Xover analysis of the vectorized capsid sequences of RS.R3, RS.R6 and RS.R11 from the neuron selection and RS.N8d from the flowthrough selection. However, unlike the Xover analysis, we did not find any aa between 537 to 608 region AAV3B/LK03 in the flowthrough or neuron selection screen derived capsids to be significantly different from input sequences.

**Figure 5.**
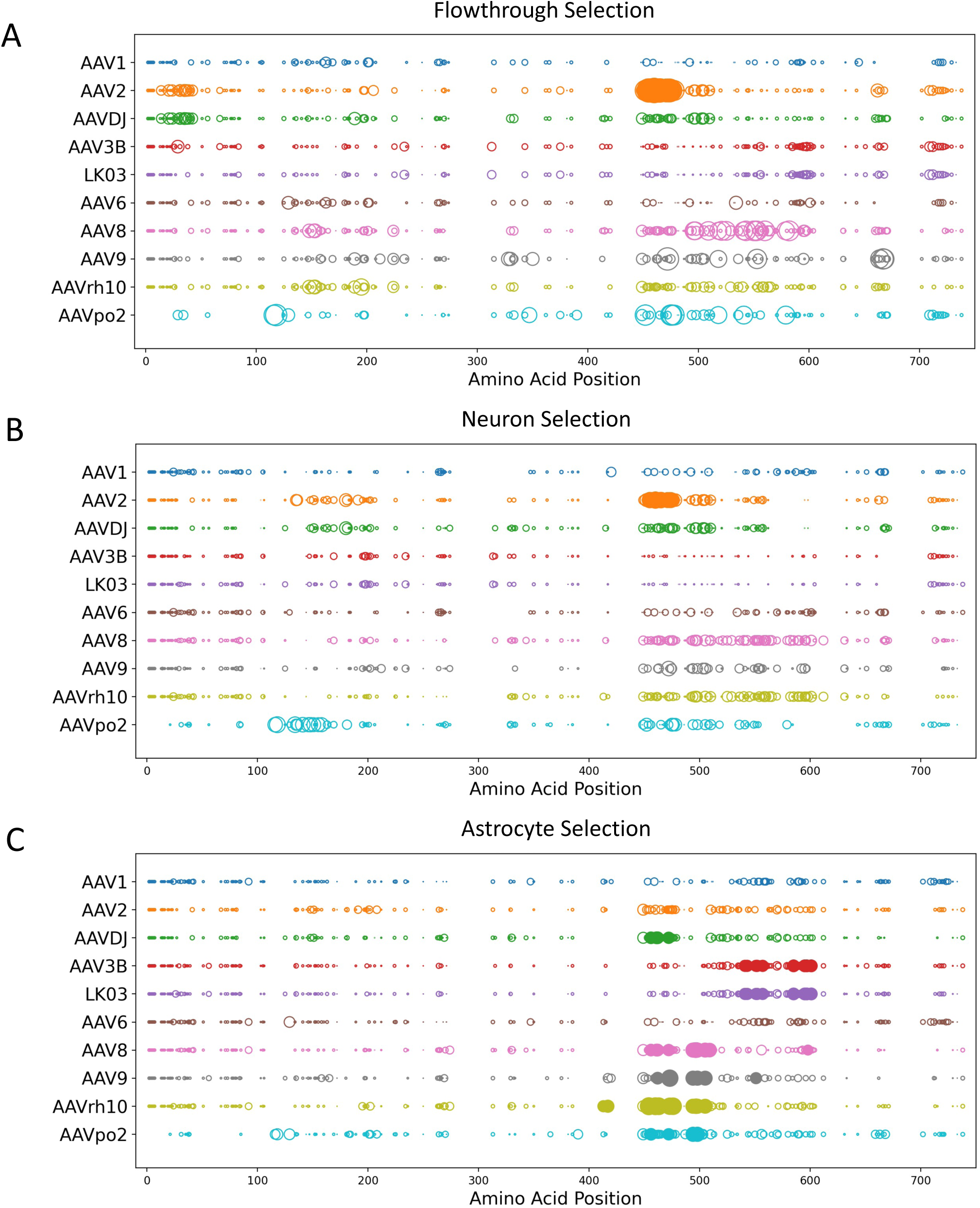
Long read sequencing analysis of AAV capsid amino acid sequences from different selections. Fold change of parental AAV contribution at each amino acid position comparing 6, 28 and 43 full capsid sequences from flowthrough (A), neuron (B) and astrocyte (C) selection, respectively, to 11613 sequences from the unselected input AAV library is plotted as circle size. Fold changes of larger than 1 is plotted, and larger circles indicate higher parental contribution at given amino acid position. Parental AAVs are depicted in different colors. Filled circles indicate statistical signifcicant (Fisher test, p <0.05) change compared to unslected input AAV, empty circles indicate non-statistical signifciant change.

Analysis of the 43 capsid sequences from the astrocyte selection identified significant contribution of aa from seven of the ten AAV parents, out of which AAV-rh10, AAV8 and AAV3B/LK03 contributed unique aa (Figure 5C, S4C, Supplemental table 2). Similar to the Xover analysis of the vectorized capsid sequences from astrocyte selection, we found AAV-rh10 contributing residues in the 413 to 496 region, with unique aa 413E, 417Q, 453S, 459G, 462Q, 475A 495L and 496S significantly enriched compared to sequences from input. We also found AAV3B/LK03 contributing to the 538 to 598 region with the exact same unique aa identified by Xover analysis (538H, 540N, 549T, 554E, 582N, 592T, 594R and 598D) showing significant enrichment compared to input. This data confirms that a combination of AAV- rh10 aa in the 413 to 496 region and AAV3B/LK03 aa in the 538 to 598 stretch may confer ability to penetrate endothelial cells and transducing astrocytes. In contrast to the Xover analysis, we also found AAV8 contributing three unique amino acids 495T, 496G and 508G. Further validation will be required to confirm whether these amino acids from AAV8 also contribute to penetrating endothelial cells and transducing astrocytes.

### siRNA screening to identify receptors that mediate transcytosis of AAV across the BBB

Transport of macromolecules from the blood to the brain across the BBB is tightly regulated by receptor-mediated transcytosis. Several receptors expressed on endothelial cells have been reported to mediate transcytosis of various types of cargo (Supplemental Table 3), and we hypothesize that these receptors may be utilized by AAVs that are efficient at crossing the BBB for transcytosis. Crossing the BBB is likely a complex process involving multiple independent or interacting pathways. Depending on the receptor engaged, endocytosed AAV capsids may be targeted for transduction of endothelial cells, transduction of astrocytes or transcytosis across the BBB. To determine which of these receptors mediate the transcytosis of AAV capsids, we used an siRNA approach to knock down the receptors in hCMEC/D3 cells grown in transwells. We then added the pool of rAAV vectors consisting of equimolar of AAV9, AAV-DJ, and representative rAAV vectors from each selection (RS.R5, RS.R6, RS.R11, RS.R18, RS.A5e and RS.A6), and tested their ability to enter or transcytose the hCMEC/D3 and primary astrocyte cell layers in the transwell system.

Knocking down of several scavenger receptors (MSR1, SCARB1 and SCARA3), insulin-like growth factor receptors (IGF1R and IGF2R), antibody transporting receptors (FCGRT and TMEM30A), as well as LEPR, LRP8, CNR1, EPHA2, PTAFR, and CCR2, resulted in reduced transcytosis of two or more rAAVs (Figure 6 and Supplemental Table 4). This observation suggests that these receptors may be important mediators of rAAV transcytosis. Knocking down of MSR1, SCARB1, SCARA3, IGF1R, IGF2R, FCGRT, LEPR, LRP8, CCR2 and MELTF also lead to increased amount of rAAV genomes in the endothelial cells and/or astrocytes, suggesting these receptors may function in the exocytosis of rAAV vectors from endothelial cells and/or directing them to astrocytes (Figure S5, S6 and Supplemental Table 4). Knocking down CNR1, EPHA2 and PTAFR did not significant change the level of rAAV genomes in the endothelial cells but reduced the levels in astrocytes for some capsid variants, suggesting that these receptors likely mediate the transfer of AAV from endothelial cells to astrocytes. Unlike the other receptors, knocking down the TMEM30A also reduced the number of rAAV genomes in endothelial cells, suggesting that this receptor may affect global endocytosis of rAAV vectors.

**Figure 6.**
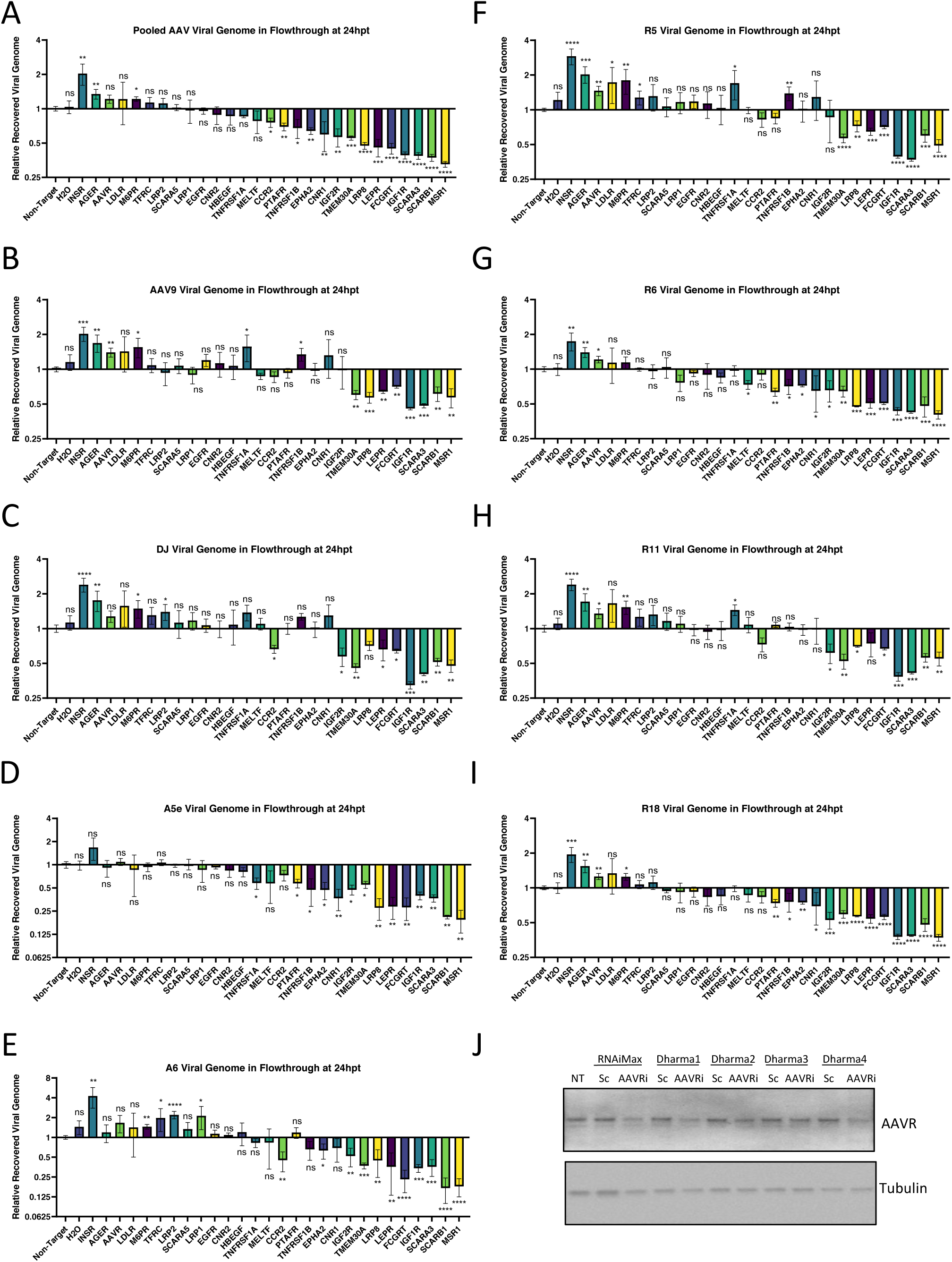
siRNA screen to identify receptors that mediate transcytosis of AAV across the BBB. A-I siRNA was used to knock down 27 receptors in hCMEC/D3 cells grown in transwells containing both hCMEC/D3 and primary astrocytes. The individually barcoded rAAV vectors of AAV9, AAV-DJ, RS.R5, RS.R6, RS.R11, RS.R18, RS.A5e and RS.A6 were pooled and passaged through these transwell BBB model with siRNA treatment. The number of vector genomes of each virus in the flowthrough of every siRNA condition were normalized to that of non-target siRNA. N = 3 for all conditions, significance determined by Student’s t test, ns, not significant; * p<0.05; ** p <0.01; *** p<0.001; **** p<0.0001. J. siRNA knock down of AAVR protein expression in hCMEC/D3 cells.

INSR, AGER, AAVR, M6PR, TFRC and LRP2 may participate in the inhibition of rAAV transcytosis across the endothelial and astrocyte cell layers, as siRNA knock down led to increased amount of two or more rAAVs in the flowthrough portion of the transwell (Figure 6 and Supplemental Table 4). Apart from AAVR, knocking down of the receptors in this category also increased the level of rAAV genomes in endothelial cells, suggesting that these receptors and the pathways they are involved in may compete for resources needed for transcytosis of rAAV (Figure S6 and Supplemental Table 4). AAVR is the receptor mediating cell entry for several AAVs including AAV9 and AAV-DJ and we show successful knock-down of AAVR as confirmed by Western blot (Figure 6J). Knocking down AAVR may have reduced the AAV directed for transduction of endothelial cells, thus increasing the amount of rAAV for transcytosis. Knock-down of INSR and M6PR also reduced the level of rAAV in astrocytes, suggesting that these receptors may play a role for AAV astrocyte transduction (Figure S5 and Supplemental Table 4).

The TNF receptors TNFRSF1A and TNFRSF1B have differential effects on different rAAV (Figure 6, S5, S6 and Supplemental Table 4). Knocking down these receptors increased the levels of AAV9 and RS.R5 in the flowthrough, but reduced their levels in endothelial cells and/or astrocytes, suggesting these receptors likely mediate their transduction of endothelial cells and astrocytes. TNFRSF1A also increased the level of RS.R11 and reduced the level of RS.A5e in the flowthrough, while TNFRSF1B reduced the levels of RS.A5e, RS.R6 and RS.R18. These results suggest that the TNF receptors may involve in multiple pathways to mediate different process for different rAAVs.

There are also several receptors that do not affect the transcytosis of rAAV across the BBB, but may impact other processes (Figure 6, S5, S6 and Supplemental Table 4). According to our data, the receptors SCARA5 and MELTF appeared to play a role in endocytosis and transduction of endothelial cells for all rAAVs tested. Receptors LDLR, CNR and HBEGF are likely involved in the transduction of astrocytes for several rAAVs, as knocking down LDLR increased vector genome copies while knocking down CNR or HBEGF reduced vector genome copies in astrocytes.

## Discussion

This current study has focused on selecting AAV vectors that efficiently cross the human BBB in an *in vitro* transwell cell culture model, and suggest this approach is an effective alternative model to identify AAV capsids with improved human CNS properties. The fact that our transwell model allowed us to predict relative enhancement or reduction in BBB penetration of a number of known capsids already tested in non-human primate pre-clinical studies and/or human trials suggests that this is an approach worth further consideration. Using this model, we have expanded the repertoire of AAV vectors that are not only efficient at transcytosing endothelial cells but also exhibit improved transduction of hiPSC-derived cortical neurons or primary astrocytes compared to the current clinical gold-standards, AAV9 and AAV-rh10. We also used this model to explore the molecular mechanisms that allow these AAV capsids to exemplify such improved performance. Specifically, using long-read high-throughput PacBio sequencing to analyze a large repertoire of capsid sequences, we were able to highlight distinctive capsid features that possibly contribute to the improved performance of our selected capsids. To elucidate some of the cellular factors that play a role in transcytosis, we knocked down a wide range of receptors known to mediate transcytosis and evaluated their effects on AAV transcytosis. We showed that several receptors may play distinct roles in targeting the virus to its destination. Overall, our work demonstrates that the transwell culture model of the human BBB is not only effective for selecting AAV capsids that may be useful for CNS based gene therapy applications but is also a valuable tool for studying the biological pathways that allow AAV to cross and target the different cell layers in the human BBB.

We opted to use the transwell culture model system of the human BBB for our studies due to its ability to overcome the limitations of using non-human models for AAV capsid selection. We also prefer this system as it is very easy to set up and is very versatile in the cell types that can be used to satisfy different selection purposes. This model has been commonly used for study of the structure and functions and BBB in supporting the understanding of how homeostasis is achieved between the transport of molecules between the blood and the brain.^16^ Previous studies have used simplified versions of the transwell BBB model to evaluate the ability of AAV in transcytosing the endothelial cell layer. In a transwell model consisting of primary human microvascular endothelial cells, Merkel et al. were the first to show that AAV9 penetrates the endothelial cell barrier more effectively than AAV2.^19^ More recently, Zhang et al. adopted the transwell BBB model consisting of a hCMEC/D3 cell barrier and showed that shuttle peptides bound to AAV8 enhanced its ability to cross the barrier.^30^

In our current study, we have taken the use of the transwell model to the next level in various ways. By incorporating primary astrocytes to the bottom of the transwell that allow direct interaction with hCEMC/D3 cells, our model not only provides a closer resemblance to the BBB, but also enabled AAV selection and evaluation in astrocytes. Furthermore, we extended our AAV selection and evaluation in hiPSC-derived cortical neurons, eliminating capsids that may have high efficiency in penetration of the cell layers but not efficient at transducing neurons. With a model of improved complexity in cell types and closer resemblance of the BBB, we evaluated the performance of 18 AAVs with different capsid sequences and performed several selection regimens using a highly complex capsid-shuffled AAV library. Our results show that the transwell BBB model consistently enriched for those capsids that confer enhanced efficiency in crossing the cell layers and in transducing various cell types. Our transwell BBB model was also able to capture capsids such as AAV-DJ, which is particularly poor at crossing the BBB but efficient at transducing human hiPSC-derived cortical neurons and astroglia cells shown here and in organoid models.^8^ Interestingly, we also found AAV-DJ highly enriched and expressed in the endothelial cells in our transwell model, suggesting it might be useful in disorders caused by dysfunctional BBB, such as 22q11.2 deletion syndrome.

The transwell BBB model remains a static model where the top compartment does not flow and thus may not fully represent the interactions AAVs will have with the endothelial cells in a microvasculature wall. For future selections and evaluations of AAVs that cross the BBB, utilization of the novel microfluidic models may prove to be more relevant.^32^ Furthermore, selection on BBB models alone does not guarantee the output AAV capsids are specific for CNS targeting. As exemplified in our 18 AAV validation assay in the transwell BBB model, AAV8 showed similar efficiency at crossing the BBB as AAV9 and AAV-rh10. Although intravascular administration of AAV8 has been previously shown to result in efficient brain cell transduction, this serotype remains a robust vector targeting the liver.^33^ In future screens, to eliminate selection of AAVs that may potentially target tissues other than the CNS, it may be beneficial to perform parallel on-chip selection schemes for various organs, such as liver, pancreas, kidney, and choose mutually exclusive AAV vectors.

As with any human cellular model, the value of the model in prediction of human application will take more effort and time to justify. Until both NHP and human clinical trials engage additional naturally occurring and chimeric AAV capsids that have shown enhancement in a model such as the one used here, the true value of the model may not be appreciated. An example is AAV-LK03, the first chimeric capsid isolated from a library screen in a humanized liver model to be used in a clinical trial.^22, 34, 35^ This capsid has been shown to result in similar if not higher levels of human factor VIII when compared with a trial utilizing AAV5 at 30 to 50x lower doses. Thus, confirming the effectiveness of the AAV capsids in crossing the BBB in NHP model and clinical trials is of crucial importance for supporting the use of the transwell *in vitro* model for prediction of capsids with improved properties for treatment of CNS diseases.

Understanding the molecular mechanism of how AAV vectors cross the BBB is also an important aspect for the development of future vectors. Previous studies have shown that transcytosis of AAV is a complex process that likely involves multiple cross-reacting pathways. Merkel et al. have shown that AAV2 tends to accumulate in oval shaped vesicular structures and nucleus, whereas AAV9 accumulates in tubular vesicular formations and basolateral plane of endothelial cells.^19^ This observation correlates well with the finding that AAV2 has a higher transduction rate for endothelial cells while AAV9 has a higher rate of penetration through endothelial barriers. Thus, we hypothesize that interaction of the AAV capsid with specific receptors at the at the endothelial cell surface may determine the destiny of this particular vector, resulting in either the transduction of the endothelial cells, the transport to astrocytes in direct contact with the endothelial cells, or the trafficking to the basolateral surface followed by dissemination to adjacent tissue.

Through Xover analysis of the vectorized AAV capsids and long read sequencing analysis of selected pools, we identified specific amino acid signatures that are associated with distinct phenotypes. In AAV capsids selected for transduction of astrocytes, the aa 413 to 496 of AAV-rh10 and amino acids 538 to 598 of AAV3B/LK03 have a significantly high frequency of occurrence. While many residues within the 413 to 496 aa region of AAV-rh10 have been suggested to confer the ability to cross BBB in previous studies ^29^, the combination with a stretch derived from AAV3B/LK03 may be crucial for the association with cellular factors that result in astrocyte transduction. The capsids selected from the flowthrough and neuron screens showed a high contribution rate of AAV2 residues from aa 451 to 470. This is an unexpected finding as no previous studies have shown these AAV2 residues confer transcytosis capabilities. Further characterization and evaluation will be needed to confirm that these aa patterns are truly responsible for the observed phenotypes. Whether these aa residues interact with specific receptors that target AAV capsids to distinct cellular pathways will be another line of study worth pursuing.

Using an siRNA approach to knock down expression of various receptors that are known to mediate transcytosis of macromolecules across the BBB, we confirmed the hypothesis that different receptors may direct AAV capsids to different destinations. Based on our data, SCARA5 and MELTF are likely involved in the transduction of endothelial cells. Receptors INSR and M6PR, LDLR, CNR and HBEGF may regulate the transport of AAV vectors to astrocytes that are in contact with endothelial cells. We also found several receptors that may enhance AAV transcytosis (MSR1, SCARB1, SCARA3, IGF1R, IGF2R, FCGRT, TMEM30A, LEPR, LRP8, CNR1, EPHA2, PTAFR, and CCR2), while others appeared negatively regulate the process (INSR, AGER, AAVR, M6PR, TFRC and LRP2). The TNF receptors TNFRSF1A and TNFRSF1B were shown to have differential effects on transcytosis depending on the AAV capsid, suggesting that these receptors may be involved in multiple pathways. Further in-depth assessment will be needed to characterize if these receptors directly interact with AAV capsids to regulate the targeting process or play an indirect role.

Taken together, our studies show that the in an *in vitro* transwell BBB cell culture model is suited for selection of AAV vectors that efficiently transcytose and transduce certain cell types in the BBB. We hope to evaluate the efficacy and safety of these vectors in the NHP model and move these capsids to clinical use in gene therapy for CNS diseases. We also elucidated molecular mechanisms that may be responsible for the improved phenotype by detailed sequencing analysis and receptor knock-down experiments. We hope that the further characterization of these mechanisms will support future endeavors in rational design of better AAV vectors for clinical applications.

## Materials and Methods

### Cell cultures

hCMEC/D3 cells (Cat #SCC066) and media (Cat #SCME004), dish coating material type I rat collagen (Cat #08-115) were purchased from MilliporeSigma, MA. Primary human astrocytes isolated from cerebral cortex (Cat # 1800), astrocyte medium (Cat # 1801), trypsin neutralization solution (Cat # 0113) and dish coating material poly-L-lysine (Cat # 0403) were purchased from ScienCell Research Laboratories, CA. All cells were maintained according to manufacturer instructions. Generation and culture of hiPSC-derived neurons and astroglia from human cortical spheroids (hCS) were described previously.^17, 26, 36^<u>_ENREF_35</u> Briefly, hiPSC derived from fibroblasts and maintained on MEFs were used to generate hCS for more than 150 days, after which they were enzymatically dissociated and immunopanned into purified neurons (Mouse anti-Thy1 (CD90), BD Biosciences, Cat. 550402) and astroglia cells (Mouse anti-HepaCAM, R&D, Cat # MAB4108) as previously described.^26^ The isolated cells were seeded on poly-D-lysine coated plates and maintained in a Neurobasal/DMEM based serum-free medium.^26^ All cell cultures were maintained in a humidified incubator at 37°C with 5% CO_2_. All human stem cell work was performed with approval from the Stanford Human Stem Cell Research Oversight (SRCO) committee.

### Culture of Transwell human BBB model

For construction of human BBB cultures in transwell, we followed a previously published protocol with some modifications.^37^ Briefly, for cultures containing hCMEC/D3 cells only, cells were seeded at 2 x 10^5^ cells/cm^2^ on the top of a 0.4 μM PET membrane (Cat # 3450 or 3470, Corning, NY) pre-coated with collagen. The 0.4 μM pore size was selected as it prevents cells from crossing between top and bottom sides, but allow AAVs of 20 nM to cross. Cultures containing both hCMEC/D3 cells and human primary astrocytes were set up as follows: astrocytes were seeded at 6 x 10^4^ cells/cm^2^ on the bottom side a 0.4 μM PET membrane pre-coated with poly-L-lysine. After 24 h of incubation, hCMEC/D3 cells were seeded at 2 x 10^5^ cells/cm^2^ on the top membrane after coating with collagen. For all transwell BBB cultures, heparin sulfate included in the hCMEC/D3 media kit was omitted as it interferes with binding of several AAV serotypes to heparin sulfate proteoglycans. Full media changes were performed every 2 to 3 days, and cultures were maintained for 11 days after seeding of hCMEC/D3 cells before use for selection of AAV for BBB penetration.

### Testing permeability in transwells using fluorophore-conjugated dextran

Transwells with hCEMC/D3 culture only or with both hCMEC/D3 and primary astrocytes were tested for permeability by adding to the top of the transwell: 70k MW Dextran-TRITC (Invitrogen, MA) (ex 555nm/em 580nm) at 1μM and 2M MW Dextran-FITC (Invitrogen, MA) (ex 494nm/em 521nm) at 5nM. Media from the bottom of the transwell were collected at 2h, 4h, 6h, and 8h after addition of the dextran. Fluorescent intensity of the collected media were measured using Tecan Infinite M1000Pro (Tecan, Switzerland).

### Production, purification and titer of AAV libraries

The generation of 18 uniquely barcoded AAV pool and 10 parent capsid-shuffled and barcoded AAV library were described previously.^18^ Briefly, the 18 AAV pool contains serotypes (AAV1, AAV2, AAV3B, AAV4, AAV5, AAV6, AAV8, AAV9.hu14, AAV12, AAV-porcine1, AAV-porcine2, AAV-bovine, AAV-goat1, AAV-mouse1, AAV-avian, AAV-rh10, AAV-DJ, and AAV-LK03), which were separately cloned into a vector containing unique barcode sequence. AAVs were pooled at equimolar prior to purification. For the 10 parent capsid-shuffled AAV library, capsid sequences from AAV1, AAV2, AAV-DJ, AAV3B, AAV-LK03,AAV6, AAV8, AAV9.hu14, AAV-porcine2, and AAV-rh10 were used as parental sequences for the shuffling and resulted fragments were cloned into a BC library vector. The 18 AAV pool and 10 parent capsid-shuffled AAV library were purified by two consecutive CsCl gradient centrifugation, and genome titers were determined by quantitative polymerase chain reaction (qPCR) using an AAV2 *REP* gene-specific primer/probe set described in Ref ^18^.

### Selection of 18 AAV pool and 10 parent capsid-shuffled AAV library in transwell BBB model

The AAV input was added to the top side of transwell BBB culture model containing both hCMEC/D3 cells and human primary astrocytes. After 24h, astrocytes and flowthrough media on the bottom of the transwell culture were collected. In a separate transwell BBB culture, hCMEC/D3 and flowthrough media were removed at 24h and Ad5 was added to the astrocytes for 72h prior to collection. The flowthrough media was also added to hiPSC-derived cortical neuron cultures with or without Ad5 and collected after 72h. DNA from input and all collected samples were tested for AAV types by NGS of barcodes previously described in Ref ^18, 20^. For the 10 parent capsid-shuffled AAV library, second round of selections were performed with three different inputs from round one. In the first, flowthrough media from round one was the input added to a new transwell BBB culture, and flowthrough media from round two was collected after 24h. In the second, astrocytes with Ad5 from round one was frozen and thaw three times, heat inactivated at 65°C and used as input on a new transwell BBB culture. Astrocytes from the new transwell was collected after 24h. For the third, hiPSC-derived cortical neurons with Ad5 from round one was frozen and thaw three times, heat inactivated at 65°C and used as input on a new transwell BBB culture. Flowthrough media from the new transwell was collected after 24h and added to fresh hiPSC-derived cortical neurons for 72h prior to collection. Two biological replicates were performed at every selection step. Barcodes that were present in both replicates and were enriched in both rounds of selection were identified.

### Recovery and evaluation of enriched AAV capsid in the transwell BBB model

Capsid sequences that have enriched barcode count in the two rounds of selection were PCR amplified using primer capF ^18^ and a primer containing the specific barcode sequence. The capsids DNA sequences recovered from the selections were analyzed by Sanger sequencing and subjected to DNA shuffling pattern analysis using the Xover 3.0 online interface ^27^. The capsids were then used to package a single-stranded rAAV vector expressing firefly luciferase under control of the CAG promoter (pAAV-CAG-FLuc, Addgene, catalog 83281). rAAVs were purified using AAVpro® Purification Kit for All Serotypes (Cat # 6666, Takara Bio USA, CA) and each of the purified rAAVs were tested in separate transwell BBB cultures with hCMEC/D3 cells and primary astrocytes at MOI of 1e4. Flowthrough media was collected at 3, 6 and 24h and astrocytes were collected at 24h. Vector genome copies in input and collected samples were determined by qPCR using a FLuc-specific primer set described in Ref ^18^.

### Evaluation of rAAVs for transduction efficiency in hiPSC-derived cortical neurons and astroglia cells

Takara kit purified rAAVs capsids packaged with pAAV-CAG-FLuc were tested in separate cultures of hiPSC-derived cortical neurons and astroglia cells at MOI of 1e2, 1e3, 1e4 and 1e5. After 72h, cells were lysed in plate and luciferase activity was measured using Luciferase 1000 Assay System (Cat # E4550, Promega, WI).

### Evaluation of rAAVs for transduction efficiency in mice

rAAVs capsids packaged with pAAV-CAG-FLuc were purified by two consecutive CsCl gradient centrifugation. Each rAAV were test in separate B6 albino mouse (B6N-*Tyrc-Brd*/BrdCrCrl, Charles River, MA) through retro-orbital injection. Brain and liver tissues were collected 21 days or 28 days after injection. Vector genome copies in the collected samples were determined by qPCR using a FLuc-specific primer set described in Ref ^18^. All mouse procedures were approved by the IACUCs of Stanford University.

### Construction and evaluation of pooled rAAVs in the transwell BBB model

A single-stranded rAAV vector expressing human frataxin (hFXN) under control of the CAG promoter (pAAV-CAG-hFXN) was generated by replacing the FLuc sequence in the pAAV-CAG-FLuc plasmid with hFXN. For generation of the barcodes, oligonucleotides with 12 random nucleotides and restriction site sequences on both 5’- and 3’- ends followed by (5’-CTT GAG AAT TCG ATA TCN NNN NNN NNN NNA AGC TTA TCG ATA ATC AAC C-3’) were annealed to an oligonucleotide containing the antisense of the 3’ restriction site (5’-GGT TGA TTA TCG ATA AGC TT-3’) and extended using Klenow Polymerase devoid of exonuclease activity (NEB). Resulted fragments were inserted into the pAAV-CAG-hFXN plasmid between the stop codon of the hFXN and poly A signal. Individual clones were sequenced to assess the barcode identity. rAAV were generated by packaging each capsid with a pAAV-CAG-hFXN carrying a unique barcode, purified using Takara kit and pooled at equimolar ratios. The pooled rAAVs were tested in the same transwell BBB culture with hCMEC/D3 cells and primary astrocytes at MOI of 1e4. Flowthrough media, hCMEC/D3 cells, astrocytes were collected at 24h. DNA from input and all collected samples, RNA from the hCMEC/D3 cells and astrocytes were tested for AAV types by NGS of barcodes previously described in Ref ^18, 20^.

### PacBio sequencing and Analysis

All samples collected from the flowthrough, neuron and astrocyte selections of the 10 parent capsid-shuffled AAV library were used to amplify ∼2.4-kb fragments containing the capsid as well as the BC sequences. Library preparation and Pacific Biosciences (PacBio) sequencing using Sequel® Sequencing Kit 3.0 (Cat # 101-613-700) for 10h movies were performed by the University of Washington PacBio Sequencing Services. Circular consensus sequence (CCS) reads were generated using the SMRT Link 6.0.0.47841 with filtering set at Minimum Number of Passes = 3 and Minimum Predicted Accuracy = 0.9. The barcodes were first extracted from the CCS reads using seeq, a DNA/RNA pattern matching algorithm (https://github.com/ezorita/seeq). The barcodes with three or more CCS reads were included in downstream bioinformatics analyses to generate consensus DNA sequence, extract the ORF of capsid sequence and translate into amino acid sequence. The translated capsid amino acid sequences along with 10 parent sequences were subjected to MAFFT v7.427, a multiple sequence alignment program for amino acid and nucleotide sequences. Based on the alignment, the amino acid contribution from each parent at every position were calculated. Barcodes that were previously identified enriched in two rounds of selection by barcode NGS sequencing were used to pulled out full length capsid sequences. The amino acid contribution from each parent at every position were averaged for capsid sequences from flowthrough, neuron or astrocyte selections. Then fold-change and statistical significance was calculated compared to capsid sequences in the input.

### siRNA transfection and evaluation of the pooled rAAVs in the transwell BBB model

Pools of 4 siRNA targeting each receptor was purchased from Dharmacon, CO. siRNA transfection was performed on both day 6 and day 8 after seeding of hCMEC/D3 cells. Lipofectamine RNAiMax (Cat # 13778075, Thermo Fisher Scientific, MA) was used to transfect 60nM of the siRNA pool on each of the two days transfection was performed. Knockdown efficiency for AAVR was assessed by Western blot using an antibody against AAVR (Cat # ab105385, Abcam, UK). The pooled rAAVs carrying uniquely barcoded CAG-hFXN were tested in the transwell BBB culture with hCMEC/D3 cells transfected with siRNA and primary astrocytes at MOI of 1e4. Flowthrough media, hCMEC/D3 cells, astrocytes were collected at 24h. DNA from input and all collected samples were tested for AAV types by NGS of barcodes.

### Statistical Analysis

Statistical calculations were conducted using Prism v8.4 or NumPy package in Python. Experimental values were assessed via 2-way ANOVA test, Student’s t-test or Fisher’s exact test. p values of less than 0.05 were considered statistically significant.

## Acknowledgments

We would like to acknowledge Shuyuan Zhang for writing the Python scripts used in sequence analysis, Se-Jin Yoon for assistance with immunopanning experiments, Xuhuai Ji from the Stanford Genomics Facility for performing the NGS and Jeff Rubin for reviewing the manuscript. RS was supported in part by a postdoctoral fellowship from Stanford Dean’s Fellowship. This work was supported by grants to MAK NIH R01AI116698. SPP is a NYSCF Robertson Investigator, a CZI Ben Barres Investigator and a CZ BioHub Investigator. RS and MAK are inventors on patents for AAV serotypes used in this paper. Stanford has submitted patents for the AAV capsids described her in which MK and RS are inventors. The contents of this publication are solely the responsibility of the authors and do not necessarily represent the official views of the various funding bodies or universities involved. Packaging plasmids for any of the new capsids described herein must be obtained through a material transfer agreement (MTA) with Stanford University.

## Author Contributions

RS and MAK designed the experiments. RS, KP, TAK, and FZ generated reagents and protocols, performed experiments and analyzed data. RS, SPP and MAK acquired funding. RS wrote the manuscript and generated the figures. All authors reviewed, edited and commented on the manuscript.

## Supplemental Tables

**Supplemental Table 1Validation of the transwell model of human BBB using 18 AAV pool.** 18 uniquely barcoded AAVs were pooled and passaged through the transwell BBB model with hCMEC/D3, primary astrocytes and hiPSC-derived cortical neurons at the MOI of 1e4, 1e5 or 1e6. The percentage of each of the 18 AAV in the input, flowthrough media, astrocytes with or without Ad5, and in hiPSC-derived cortical neurons with or without Ad5 is shown as well as fold-change compared to input. Red indicates increase and blue indicates decrease.

**Supplemental Table 2 Long read sequencing analysis of AAV capsid amino acid sequences from different selections.** Each tab is of a different selection, “-all” shows all amino acid positions, “-significant” shows only amino acid positions where one or more AAV parent has statistical significant contribution. Red indicates of statistical significant contribution, green indicates the amino acid is unique to the parent (exceptions for AAV2 and AAV-DJ as well as AAV3B and AAV-LK03, as these two pairs of AAV capsids share a significant number of amino acids).

**Supplemental Table 3 Receptors reported to mediate transcytosis of various types of cargo across the BBB.**

**Supplemental Table 4 siRNA screening to identify receptors that mediate transcytosis of AAV across the BBB.** siRNA was used to knock down 27 receptors in hCMEC/D3 cells grown in transwells containing both hCMEC/D3 and primary astrocytes. The individually barcoded rAAV vectors of AAV9, AAV-DJ, RS.R5, RS.R6, RS.R11, RS.R18, RS.A5e and RS.A6 were pooled and passaged through these transwell BBB model with siRNA treatment. The number of vector genomes of each virus in the flowthrough, astrocytes and hCMEC/D3 cells of every siRNA condition were normalized to that of non-target siRNA. N = 3 for all conditions, significance determined by Student’s t test, red indicates significantly increased, blue indicates significantly decreased.

**Figure S1.**
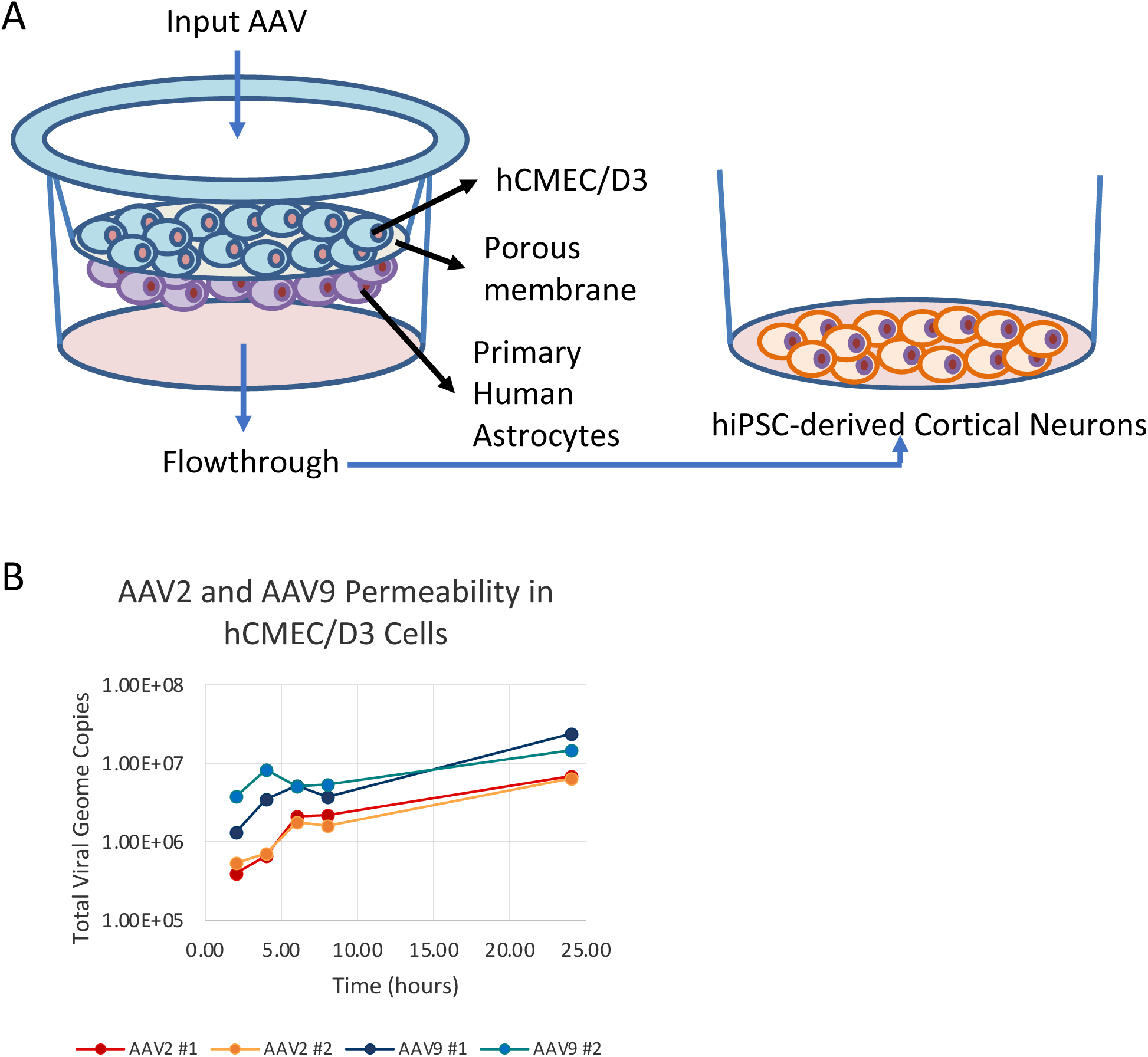
Establish and validate a transwell model of human BBB for selection of AAVs. A. Schematic drawing of the transwell BBB model where hCMEC/D3 is cultured on top of a porous membrane while human primary astrocytes are cultured on the bottom. When testing AAV transduction and penetration, input is added to the top of hCMEC/D3 cells, and samples are collected in astrocytes and flowthrough media. The AAVs in the flowthrough media can also be tested on cultured hiPSC-derived cortical neurons. B. Permeability of AAV2 and AAV9 were tested in 6-well transwells seeded with hCMEC/D3 cells at an MOI of 1e4. Total viral genome copy numbers in the flowthrough media were graphed.

**Figure S2.**
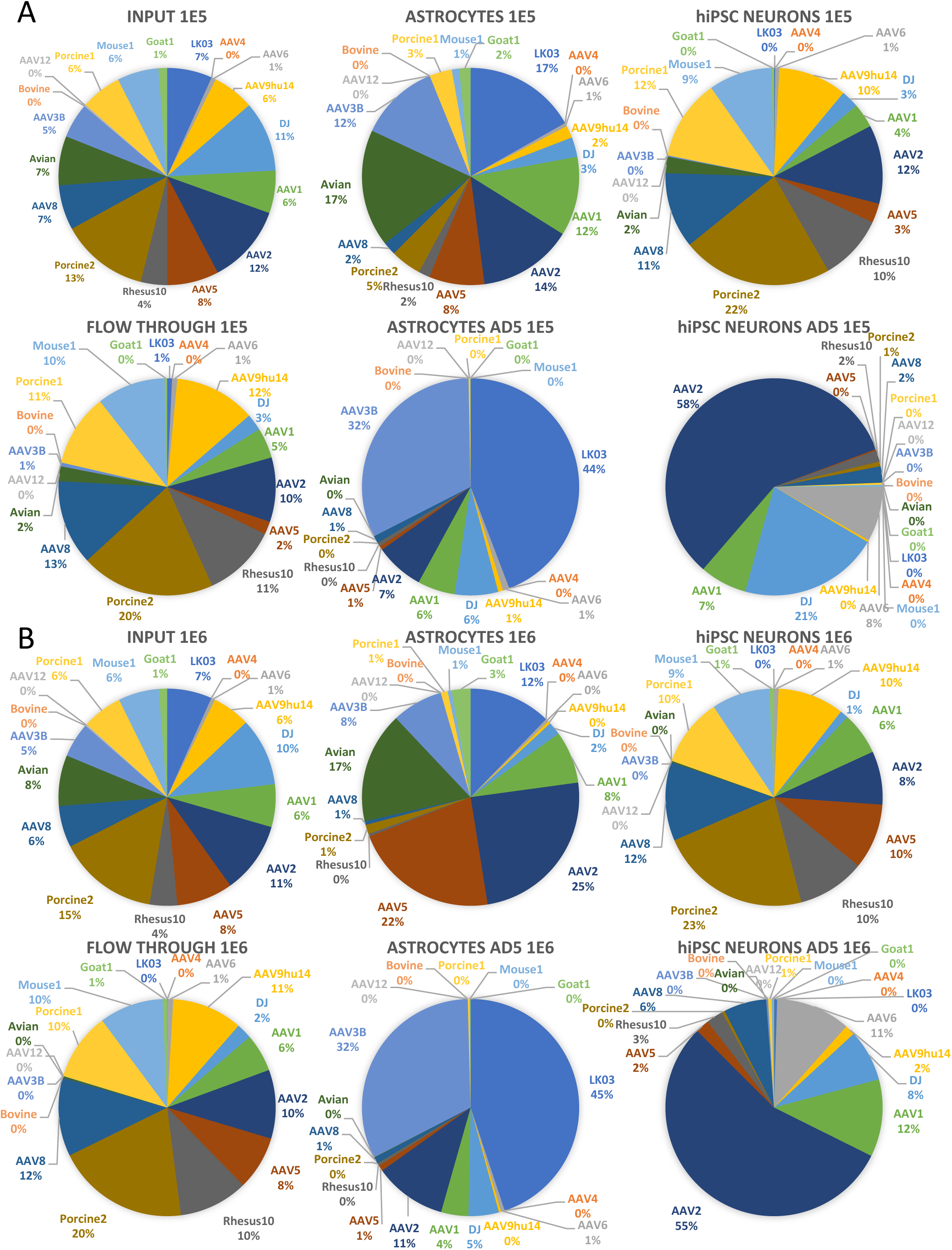
Validation of the transwell model of human BBB using 18 AAV pool. 18 uniquely barcoded AAVs were pooled and passaged through the transwell BBB model with hCMEC/D3, primary human astrocytes and hiPSC-derived cortical neurons with MOI of 1e5 (A) or 1e6 (B). The percentage of each of the 18 AAV is graphed in the input, flowthrough media, astrocytes with or without Ad5, and in hiPSC-derived cortical neurons with or without Ad5.

**Figure S3.**
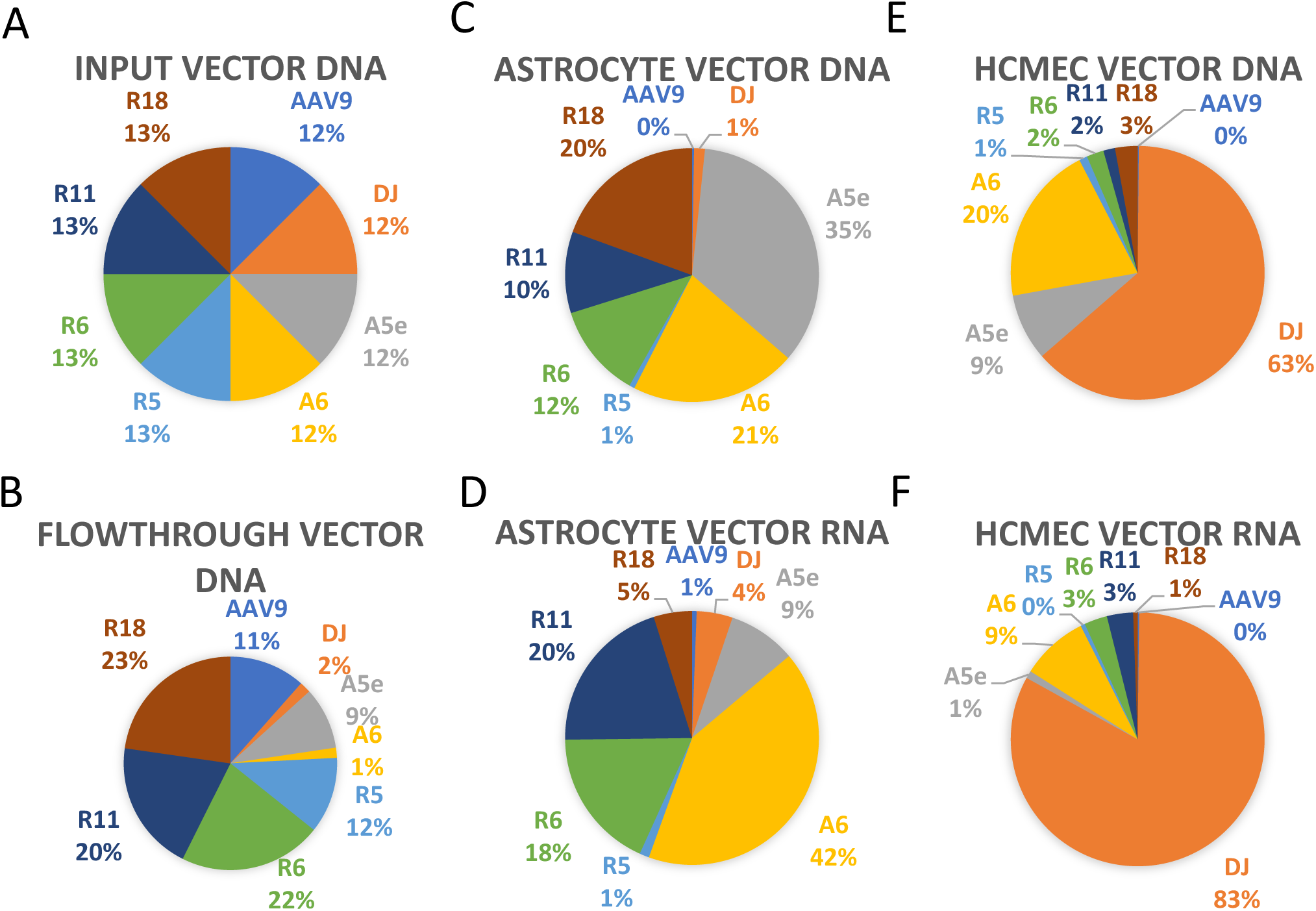
Pooled approach to validate selected rAAV vectors in the transwell BBB model. Representative rAAV vectors from each selection (RS.R5, RS.R6, RS.R11, RS.R18, RS.A5e and RS.A6) along with controls AAV9 and AAV-DJ were each package with a CAG-frataxin expression construct containing a unique barcode sequence. The rAAV vectors were pooled and passaged through the transwell BBB model with hCMEC/D3 and primary astrocytes with MOI of 1e4. The percentage of vector genome from each of the 8 AAVs were graphed in the input (A), flowthrough media (B), astrocytes (C) and hCMEC/D3 (E). The percentage of vector RNA expression from each of the 8 AAVs were graphed in astrocytes (D) and hCMEC/D3 (F).

**Figure S4.**
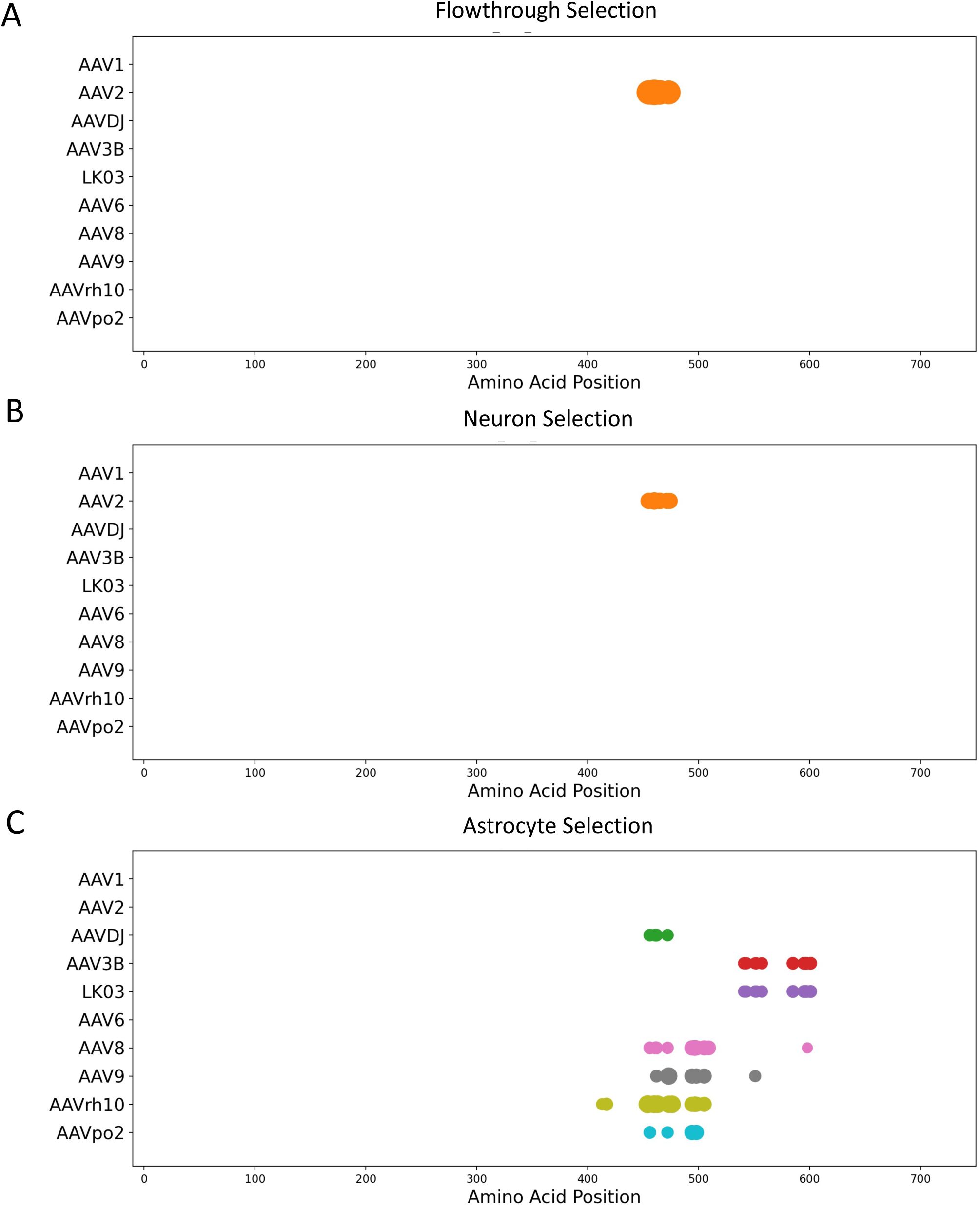
Long read sequencing analysis of AAV capsid amino acid sequences from different selections (Significan only). Fold change of parental AAV contribution at each amino acid position comparing 6, 28 and 43 full capsid sequences from flowthrough (A), neuron (B) and astrocyte (C) selection, respectively, to 11613 sequences from the unselected input AAV library is ploted as circle size. Fold changes of larger than 1 and statistically signifcicant (Fisher test, p <0.05) are plotted, and larger circles indicate higher parental contribution at given amino acid position. Parental AAVs are depicted in different colors.

**Figure S5.**
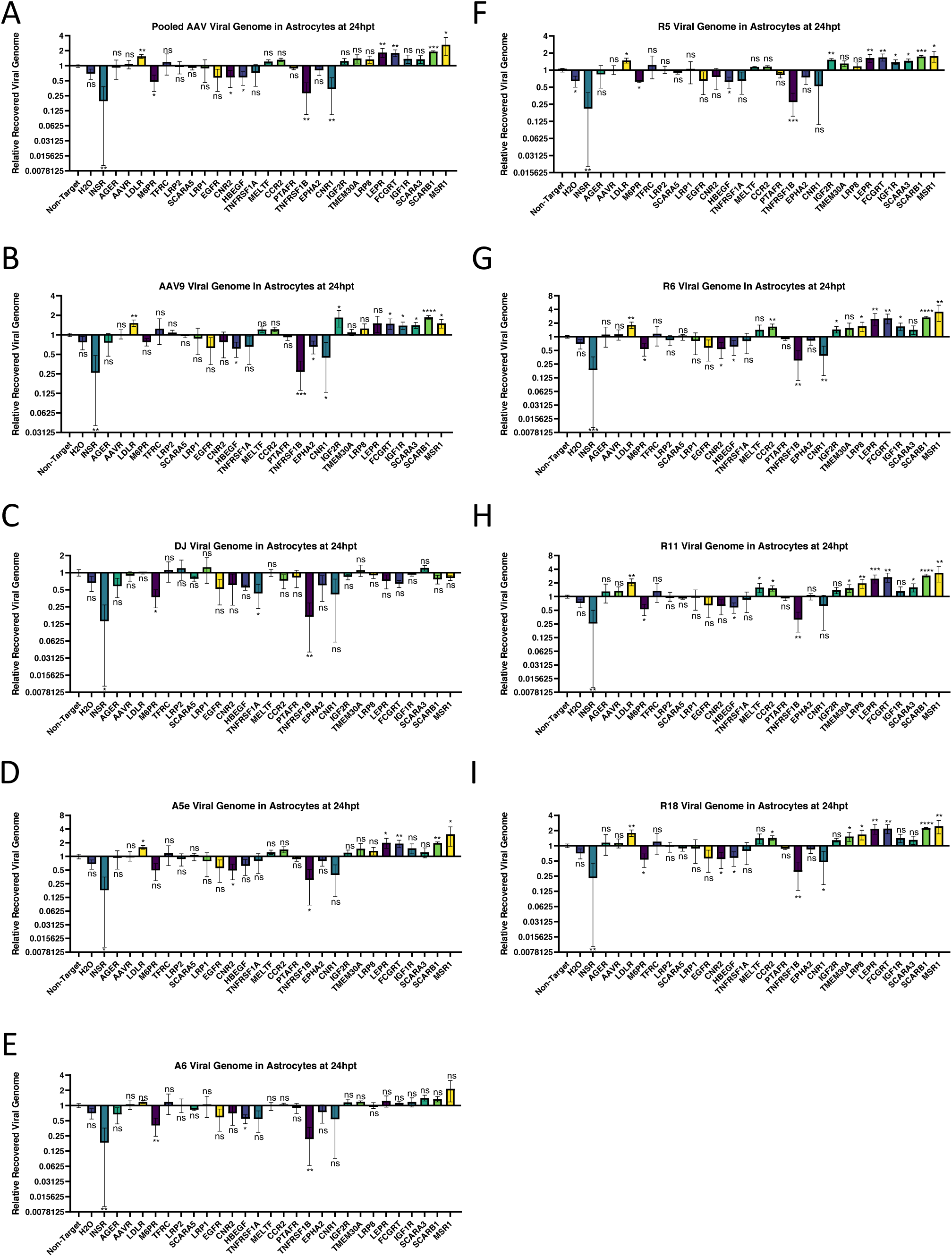
siRNA screen to identify receptors that mediate transcytosis of AAV across the hCMEC/D3 cells and transduce primary astrocytes. siRNA was used to knock down 27 receptors in hCMEC/D3 cells grown in transwells containing both hCMEC/D3 and primary astrocytes. The individually barcoded rAAV vectors of AAV9, AAV-DJ, RS.R5, RS.R6, RS.R11, RS.R18, RS.A5e and RS.A6 were pooled and passaged through these transwell BBB model with siRNA treatment. The number of vector genomes of each virus in the astrocytes of every siRNA condition were normalized to that of non-target siRNA. N = 3 for all conditions, significance determined by Student’s t test, ns, not significant; * p<0.05; ** p <0.01; *** p<0.001; **** p<0.0001.

**Figure S6.**
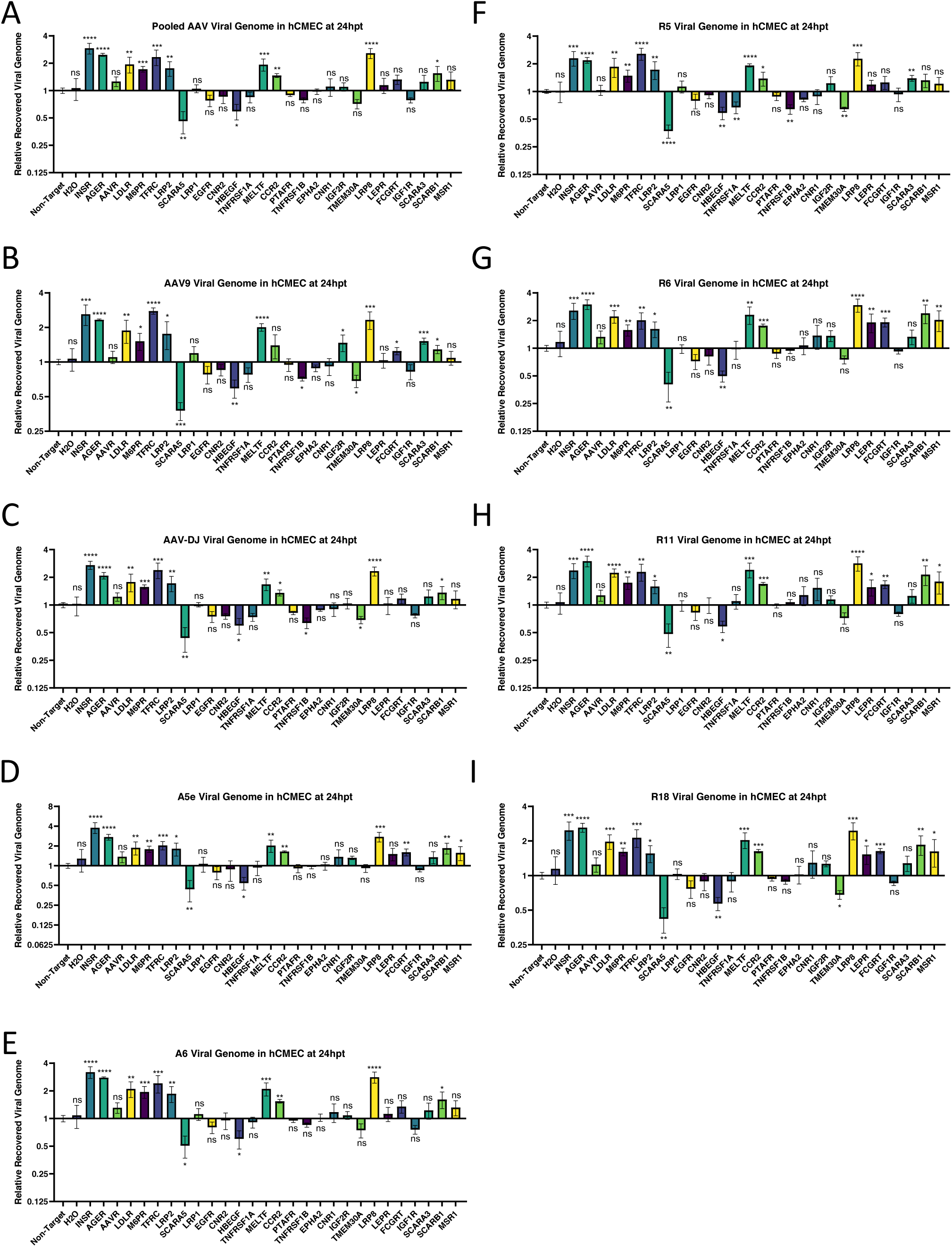
siRNA screen to identify receptors that media transduction of AAV in hCMEC/D3 cells. siRNA was used to knock down 27 receptors in hCMEC/D3 cells grown in transwells containing both hCMEC/D3 and primary astrocytes. The individually barcoded rAAV vectors of AAV9, AAV-DJ, RS.R5, RS.R6, RS.R11, RS.R18, RS.A5e and RS.A6 were pooled and passaged through these transwell BBB model with siRNA treatment. The number of vector genomes of each virus in the hCMEC/D3 cells of every siRNA condition were normalized to that of non-target siRNA. N = 3 for all conditions, significance determined by Student’s t test, ns, not significant; * p<0.05; ** p <0.01; *** p<0.001; **** p<0.0001.

## Notes

### Competing Interest Statement

Stanford has submitted patents for the AAV capsids described here in which MK and RS are inventors.

