## Supplemental Table 3 for "Selection of rAAV Vectors that Cross the Human Blood-Brain Barrier and Target the Central Nervous System Using a Transwell Model"

| Supplemental Table 3 | | | |
| --- | --- | --- | --- |
| Receptor | Symbol | Ligand | Reference |
| Transferrin Receptor | TFRC | Transferrin | ^1, 2^ |
| Melanotransferrin | MELTF | melanoransferrin | ^3^ |
| Low density lipoprotein receptor | LDLR | ApoE3, LDL | ^4^ |
| low density lipoprotein receptor related protein 1 | LRP1 | Lipoproteins, Amyloid-B, a-2-Macrglobulin, Melanotransferrin, ApoE, IgG | ^3, 5-7^ |
| low density lipoprotein receptor related protein 2 | LRP2 | Lipoproteins, Amyloid-B, a-2-Macrglobulin, Melanotransferrin, ApoE | ^8^ |
| Apolipoprotein E receptor 2 | LRP8 | Lipoproteins | ^9^ |
| advanced glycosylation end-product specific receptor | AGER | Amyloid-B, S-100, amphoterin, Glycosylated proteins | ^10^ |
| Insulin Receptor | InsR | Insulin | ^11, 12^ |
| Transmembrane protein 30A | TMEM30A | FC5 | ^13^ |
| Neonatal Fc Receptor | FCGRT | IgG | ^14^ |
| Leptin Receptor | LEPR | Leptin | ^15, 16^ |
| insulin like growth factor 1 receptor | IGF1R | IGF | ^9, 17^ |
| insulin like growth factor 2 receptor | IGF2R | arylsulfatase A, HIV | ^18-20^ |
| macrophage scavenger receptor 1 | MSR1 | Apolipoprotein A | ^9^ |
| scavenger receptor class B member 1 | SCARB1 | HDL, Apolipoprotein A, Angiopep-2 | ^21, 22^ |
| scavenger receptor class A member 3 | SCARA3 | Angiopep-2 | ^21^ |
| scavenger receptor class A member 5 | SCARA5 | Angiopep-2 | ^21^ |
| TNF receptor superfamily member 1A | TNFRSF1A | TNFa | ^9^ |
| TNF receptor superfamily member 1B | TNFRSF1B | TNFa | ^23^ |
| epidermal growth factor receptor | EGFR | EGF | ^9^ |
| heparin binding EGF like growth factor | HBEGF | Diphtheria toxin | ^24^ |
| EPH receptor A2 | EPHA2 | Cryptococcus neoformans | ^25^ |
| cannabinoid receptor 1 | CNR1 | Amyloid-B | ^26^ |
| cannabinoid receptor 2 | CNR2 | Amyloid-B | ^26^ |
| mannose-6-phosphate receptor | M6PR | arylsulfatase A | ^9^ |
| C-C motif chemokine receptor 2 | CCR2 | CCL2 | ^27^ |
| platelet activating factor receptor | PTAFR | Pneumococcal | ^28^ |
